# Whole-genome-based *Helicobacter pylori* geographic surveillance: a visualized and expandable webtool

**DOI:** 10.1101/2021.03.29.437451

**Authors:** Xiaosen Jiang, Zheng Xu, Tongda Zhang, Yuan Li, Wei Li, Hongdong Tan

## Abstract

*Helicobacter pylori* exhibits specific geographic distributions that related to the clinical outcomes. Despite the high infection rate of *H. pylori* throughout the world, the genetic epidemiology surveillance of *H. pylori* still needs to be improved. Here, we used single nucleotide polymorphisms (SNPs) profiling approach based on whole genome sequencing (WGS) that facilitates genomic population analyses of *H. pylori* and encourages the dissemination of microbial genotyping strategies worldwide. A total number of 1,211 public *H. pylori* genomes were downloaded and used to construct the typing tool, named as HPTT (*H. pylori* Typing Tool). Combined with the metadata, we developed two levels of genomic typing, including a continent scale and a country scale that nested in the continent scale. Results showed that Asia was the largest isolates source in our dataset, while isolates from Europe and Oceania were comparatively more widespread. More specifically, Switzerland and Australia are the main source of widespread isolates in their corresponding continents. To integrate all the typing information and enable researchers to compare their own dataset against the existing global database in an easy and rapid way, a user-friendly website (https://db.cngb.org/HPTT/) was developed with both genomic typing tool and visualization tool. To further confirm the validity of the website, ten newly assembled genomes were downloaded and tested precisely located on the branch as we expected. In summary, *H. pylori* typing tool (HPTT) is a novel genomic epidemiological tool that can achieve high resolution analysis of genomic typing and visualizing simultaneously, providing insights into the genetic population structure analysis, evolution analysis and epidemiological surveillance of *H. pylori*.

## Introduction

*Helicobacter pylori* is one of the most sophisticated colonizers in the world that infects more than half of world’s population ranged from infants to elders (Suerbaum and Michetti 2002). It is a Gram-negative bacterium that normally colonises at the gastric mucosa of human with about 10% infection result in diseases. The typical diseases were reported as gastritis, peptic ulcer, mucosa-associated lymphoid tissue (MALT) lymphoma and gastric cancer (Ernst and Gold 2000). Globally speaking, the risks of disease and the incidence and mortality of the gastric cancer were geographically different (Group 1993).

*H. pylori* displays a distinguished mutation rate among bacterial pathogens due to the lack of genes that initiates classical methyl-directed mismatch repair (MMR) (Alm, et al. 1999). The high mutation and recombination rate made *H. pylori* genomes with enormous plasticity, facilitating this pathogen perfectly adapted to its host (Kang and Blaser 2006; Didelot, et al. 2013). It has been reported that *H. pylori* in chronic infection could be taken place through vertical and familial transmission (Agnew and Koella 1997; Messenger, et al. 1999). In within-host evolution, the mutation rate could reach ~ 30 single nucleotide polymorphisms (SNPs) per genome per year (Kennemann, et al. 2011), comparing to *Escherichia coli* at ~ 1 SNP per genome per year (Reeves, et al. 2011). With the occurrence of large recombination events, a simple and efficient way to define the geographical pattern and epidemiological surveillance of *H. pylori* is crucially needed (Yamaoka 2009; Jolley, et al. 2018).

Among all genetic typing methods recorded in previous studies (Salama, et al. 2007; Yamaoka 2009), seven-gene multi-locus sequence typing (MLST) for *H. pylori* is a current popular tool due to its simple and rapid typing strategy. The 7-gene MLST covers genes including *atpA*, *efp*, *mutY*, *ppa*, *trpC*, *urel*, *yphC* that categorize *H. pylori* into different sequence types (STs) (Achtman, et al. 1999). This 7-gene MLST typing method enables regional specific recognition based on defined STs, through which geographical pattern may be linked with the different risk of clinical disease. For example, non-African and African lineage could be in association with different risk of gastric disease (Campbell, et al. 2001). However, the resolution of seven-gene MLST was still low, which limited us to trace the epidemiological origins of *H. pylori* strains. For users, submitting the microbial genomes is essential to get the allele number before getting the typing results. The seven-gene genotypes of *H. pylori* is diverse due to the high variability of *H. pylori* genomes, which hinders the recognition of patterns directly from the 4-digit code in 7-gene MLST. In addition, there is no information of geographical patterns or visualization tool for seven-gene MLST, thus such related geographic patterns were hard to find when a new ST was found.

Here, we describe a *H. pylori* genomic typing tool, HPTT (*H. pylori* Typing Tool) using the SNP profiling based on whole-genome sequencing data. In addition to the genomic typing, HPTT also provides a phylogenetic and geographic visualization tool based on the Nextstrain framework (Hadfield, et al. 2018). This tool allows users to upload *H. Pylori* WGS data for genomic typing and uncover the possible transmission events of *H. Pylori.* It is believed that this tool can not only improve the genome typing resolutions, but also predict the origin of the epidemic *H. pylori* isolates, enabling the global surveillance of *H. pylori*.

## Methods

### *H. pylori* genomes downloaded and filtered in this study

A total number of 1,654 assembled *H. pylori* genomes were downloaded from NCBI RefSeq database (genomes available as of 4th May 2020) using ncbi-genome-download tool (version 0.2.12). The corresponding metadata of assembled genomes was searched by function using Entrez Direct (version 10.9) (Kans 2020). By metadata filtering, 1,211 genomes were selected with sample collection location available (**Table 1**). All genomes were scanned by mlst (version 2.11) with the library of MLST updated on 31 December 2020 (Jolley and Maiden 2010).

**Table 1.**
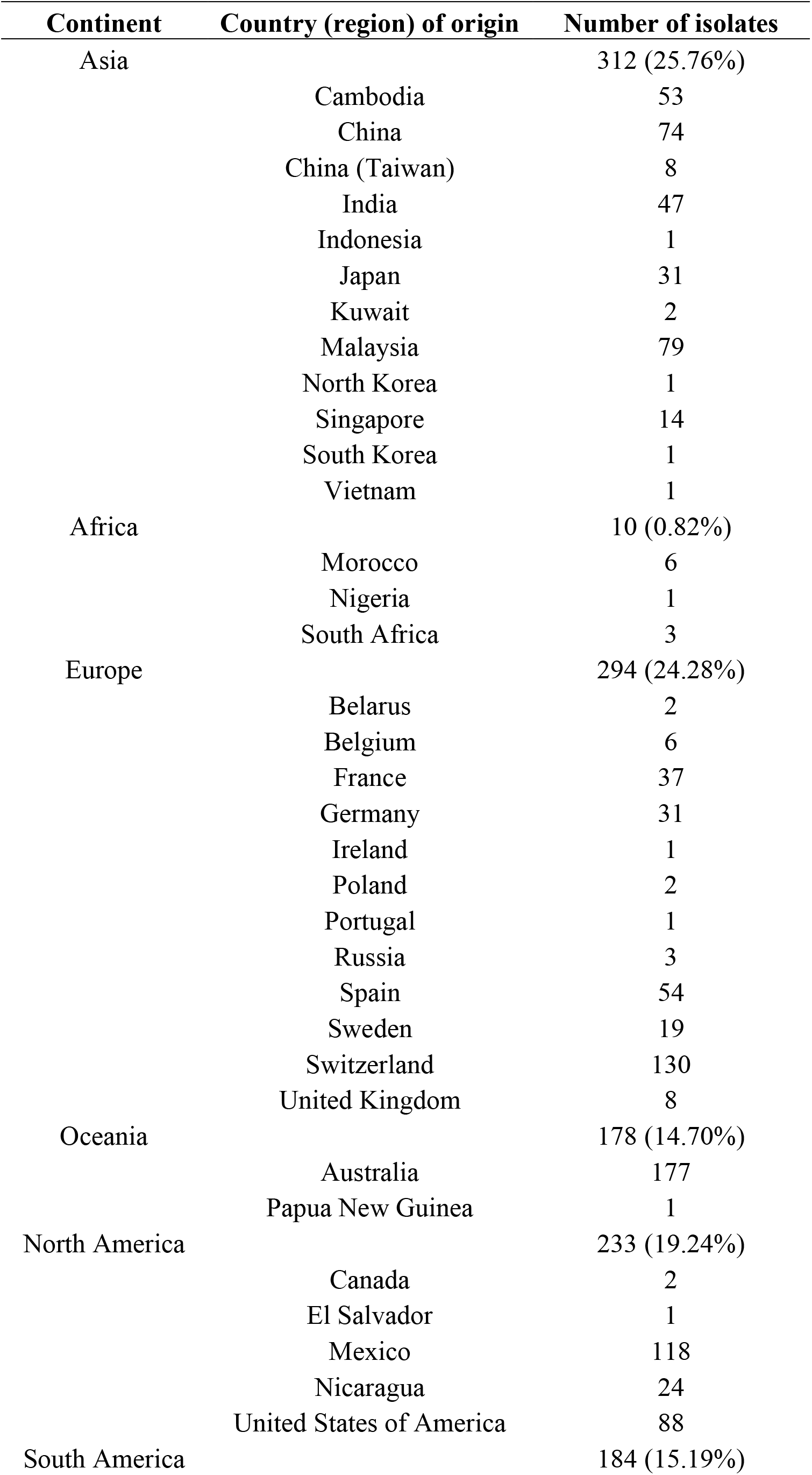
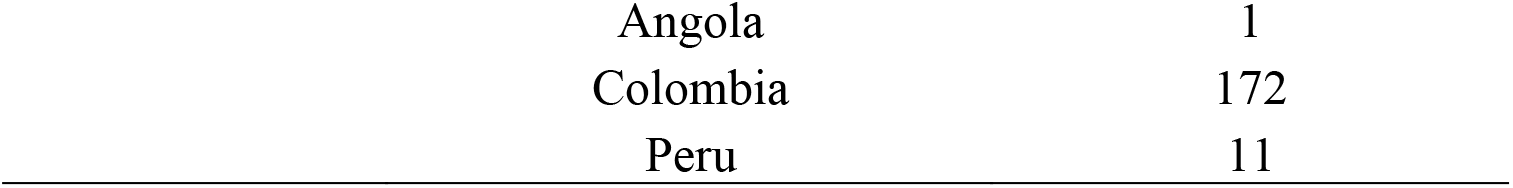
Summary of 1,211 *H. pylor* i genomes.

### SNP analysis

The 1,211 assembled genomes were mapped to the reference genome *H. pylori* 26695 (GenBank: AE000511.1) (Tomb, et al. 1997) using MUMmer (version 3.23) (Kurtz, et al. 2004). SNPs were filtered with a minimum mapping quality cutoff at 0.90 across 1,211 assembled *H. pylori* genomes. 6,129 SNPs were found, and a SNP profile of *H. Pylori* is established for the corresponding isolates.

### Phylogenetic analysis

The maximum likelihood (ML) phylogenetic tree was constructed by iqtree (version 2.0.3) (Nguyen, et al. 2015) based on 6,129 SNPs alignments of all 1,211 isolates. The reference genome *H. pylori* 26695 was used as outgroup. The tree was generalized by the Gamma distribution to model site-specific rate variation (the GTR model). Bootstrap pseudo-analyses of the alignment were set at >= 1000. All ML trees were visualized and annotated using Figtree (version 1.4.4). The minimum spanning tree was constructed by the GrapeTree (v1.5.0) (Zhou, et al. 2018).

### Geographic typing system

Based on the phylogenetic tree, two levels of geographic group were defined, including the first level defined at the continent scale and the second level defined as country specific scale. In the first level of genotyping, lineages that carrying more than seven isolates and > 75% isolates sourced from one major continent were defined as a continent specific group or clade. A mixed continent group was defined when there was no major continent identified with isolates at >75%. In the second level, lineages carrying more than one isolate and > 75% isolates sourced from one major country were defined as a country specific group or subclade. In addition, a mixed group was also defined at level two when there were more than two isolates and not a major country identified with isolates at > 75%. The association of the genomic lineage of *H. Pylori* with the geographic origin of isolates fully sequenced provide a map to allow us tracing both the origin and evolution path of a detected or sequenced *H. Pylori* genome.

### Establishment of *H. pylori* database

The HPTT website was established based on two modules: 1) The genomic-geographical typing tool of *H. pylori* isolates and 2) a visualization tool of both the genomic and geographic typing results. The online typing tool was written in PHP, Javascript, css, and html. The online visualization service was performed based on the CodeIgniter framework (https://www.codeigniter.com/), tree visualization was analysed by the augur (https://github.com/nextstrain/augur) bioinformatics tool and the auspice (https://github.com/nextstrain/auspice) visualization tool imbedded in the Nextstrain (Hadfield, et al. 2018) open source project. The *H. pylori* database was stored in a Mysql database.

## Results

### Definition of two levels of geographic genotypes for *H. pylori*

A total of 1,211 assembled genomes with available geographic information from NCBI RefSeq database were downloaded and analyzed for establishing the *H. pylori* genotyping database (**Supplementary Table 1**). All assembly genomes were mapped to the reference genome *H. pylori* 26695. Based on the maximum likelihood tree, while 6,129 SNPs were defined for further genomic typing. In terms of geographic information, 1,112 isolates were grouped at two levels, including 37 continent-level groups (**Figure 1A**) and/or 236 country-level groups (**Figure 1C**). The median pairwise distances between isolates were found as follows: 319 SNPs within continent clades and 1,493 SNPs within country subclades. We labelled these continent clades and country subclades using a structured hierarchical nomenclature system similar to that used for *M. tuberculosis* (Coll, et al. 2014). For instance, region 1 clade (G1) is subdivided into country subclades G1.C1 and G1.C2.

**Figure 1.**
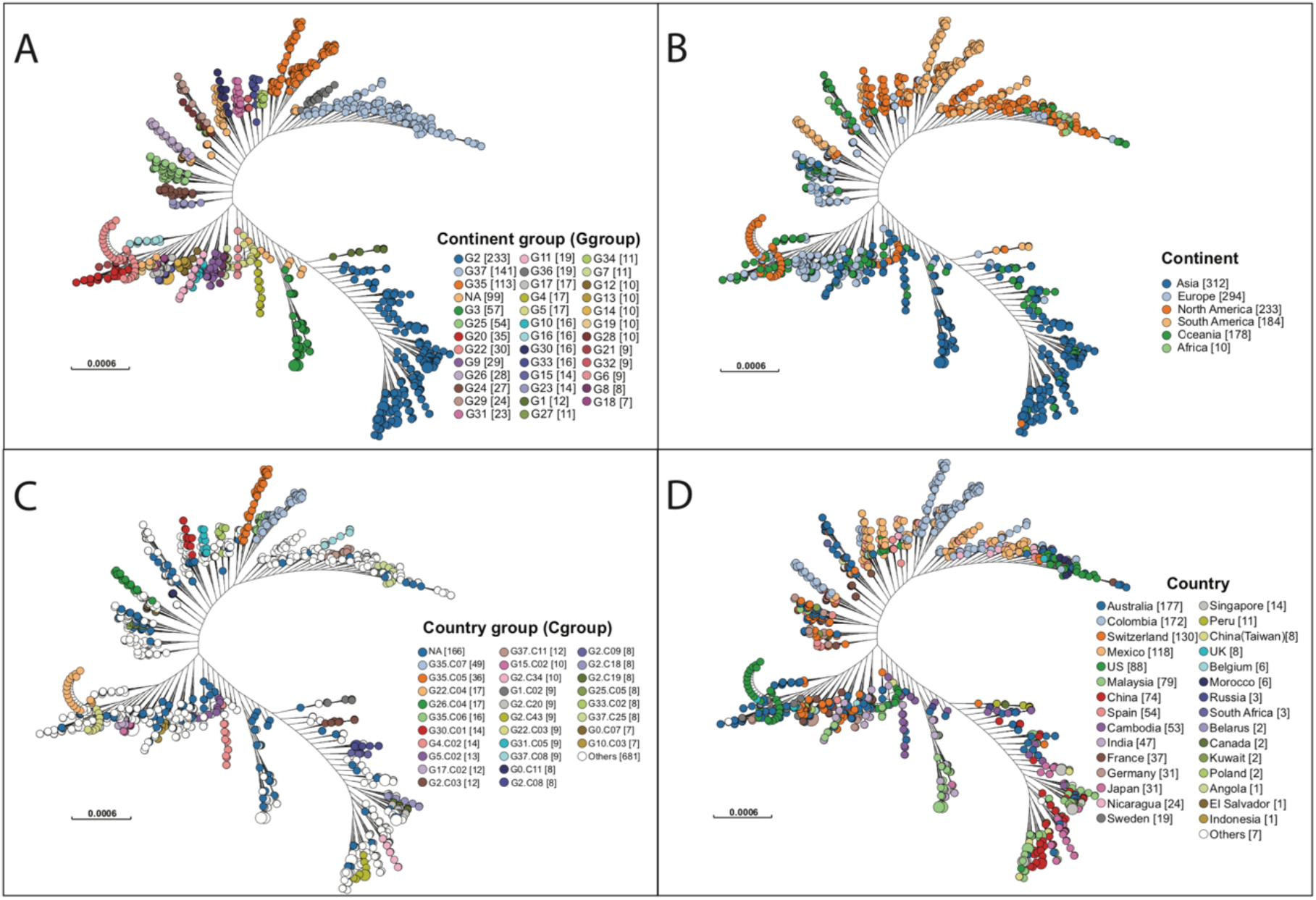
Two Clades of geographic typing based on the WGS. The HPTT enrolled 1,211 *H. pylori* genomes downloaded from NCBI. The clade nodes in each figure are corresponding to A) G groups for continent level of typing, B) the continent that isolate collected from, C) C groups for country level of typing, D) the country that isolates collected from. Numbers in parenthesis refer to the number of isolates in each genogroups.

### A continent level genomic typing for *H. pylori*

A total number of 37 continent level of groups (n=1,112) were defined, including 25 continent specific groups and the 12 mixed continent groups (**Figure 1A&B**). Isolates across the tree did not fall into the continent group but can be defined as country group were named as G0 (n=74). Isolates across the tree neither fall into the country group nor continent group was defined as non-grouped (n=25).

There were five continent specific groups contained more than 75% Asian isolates, supporting Asia to be the continent with the largest isolate source (n=319, 26.34%) (**Figure 2A**). North America was found to be the second largest group of isolate pool which consisted six continent specific groups (n=132, 10.90%). Although less isolates were found sourced from Europe (n=109, 9.00%), these isolates were distributed in nine continent specific groups. Two groups (G16 & G29) of isolates were found as Oceania specific group (n=39, 3.22%) and three groups (G1 & G26 & G35) were found as South America specific groups (n=109, 9.00%). In addition, the 12 mixed groups of isolates contained 226 isolates (18.66%). Among all G level groups, G2 was the largest continent specific group (n=223) that mainly contained isolates from Asia (193/223, 82.83%), while G35 was the second largest continent specific group (n=109) that mainly contained isolates from South America (99/109, 87.61%). Apart from all the continent groups above, there was no Africa specific group found, but only with isolates collected from Africa defined in G28 (n=2), G37 (n=7) and G29 (n=1) (**Figure 2**).

**Figure 2.**
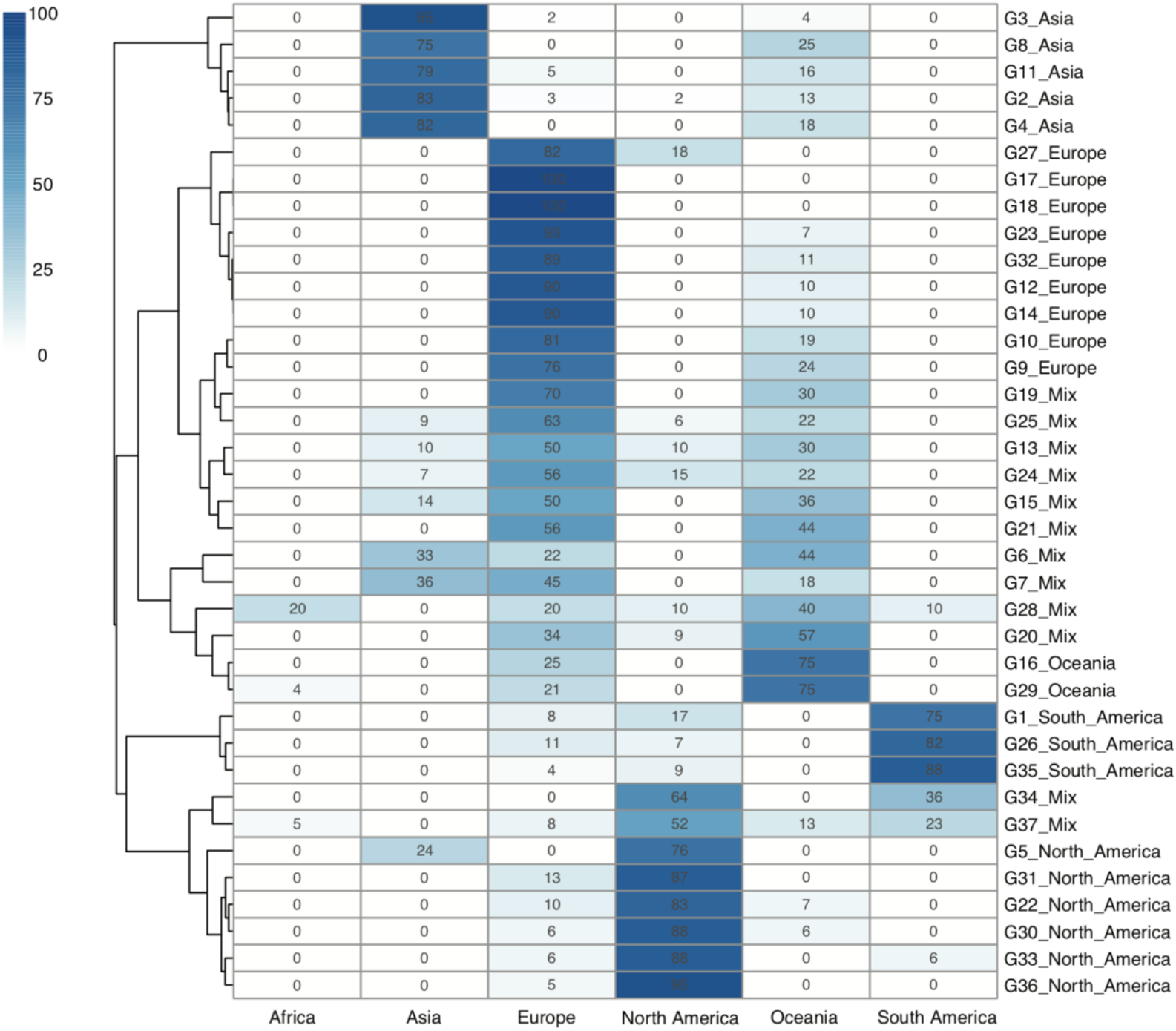
Geographical clustering of *H. pylori* continent clades. The number in each cube represents the percentage of unique isolates sourced from each of the continents. A total number of 37 continent level of groups were defined. The deeper the colour, the higher the percentage of the isolates in that continent level of clade groups. Also, a phylogenetic tree is shown in the left side of the table. The background information of isolates is provided in Supplementary Table 1.

Although the continent specific groups did not 100% stick to one continent in our typing system, the transmission events were still possible to trace. While most of the Asian isolates fell into the Asia groups, a small proportion of the Asian isolates belonged to the mixed groups. Similarly, most of the isolates sourced from North America and South America fell in their own region groups, while a minority of the isolates were in the mixed groups. Interestingly, isolates from Oceania and Europe could be found across all 12 mixed continent groups, reflecting that *H. pylori* isolates from these two continents were relatively wide spread of across the globe.

### The nested country level genomic typing for *H. pylori*

A total number of 859 isolates were grouped into 216 geographic patterns at country level, which was predominant in 29 countries across six continents (**Figure 3**). Among these 29 countries encompassing 216 groups, 20 countries found in 168 groups were defined as country-specific groups, while the rest 9 countries were scattered in the rest 48 country-level mixed groups.

**Figure 3.**
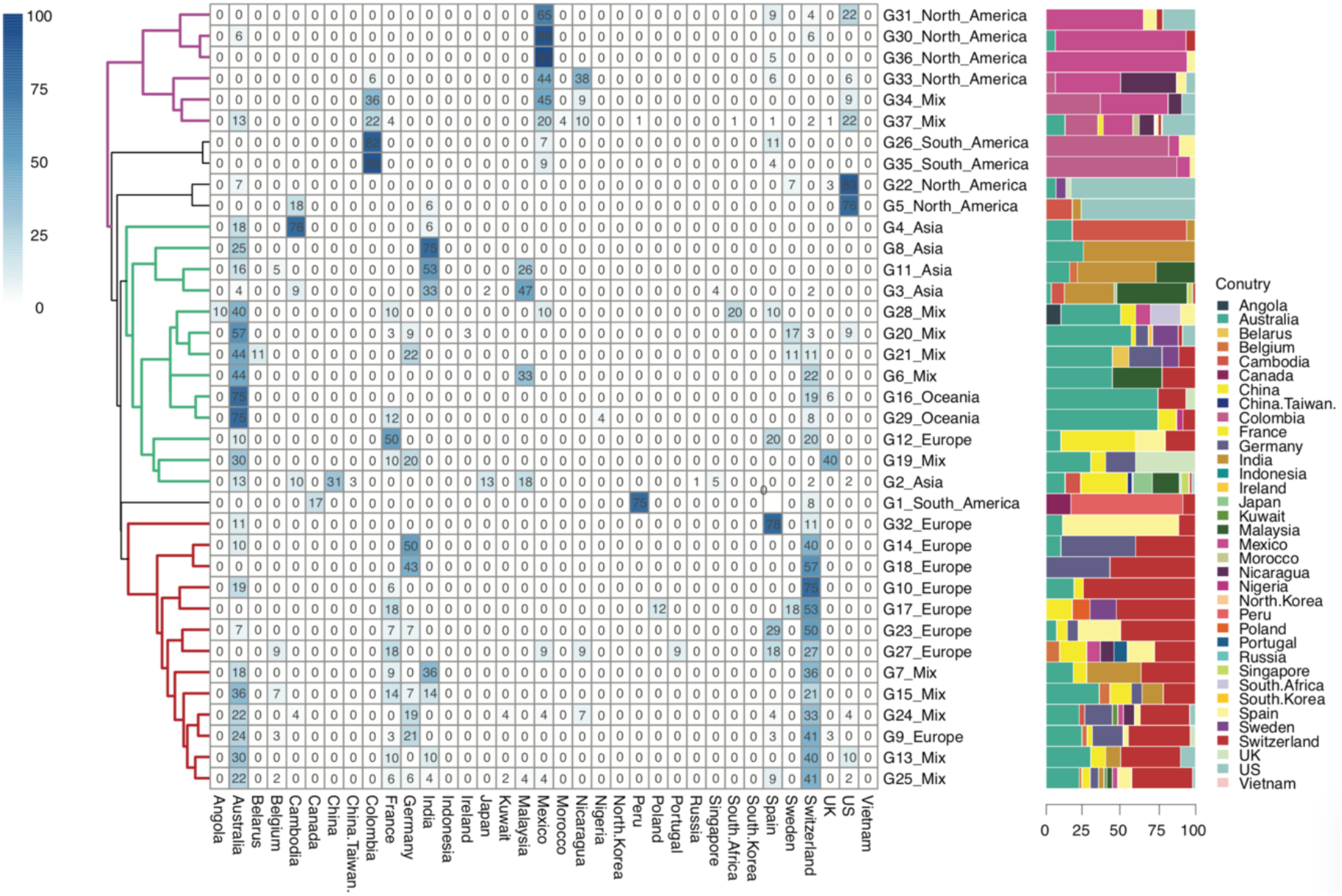
Geographical clustering of *H. pylori* country subclades. The number in each cube represents the percentage of unique isolates sourced from each of the country in that continent groups. A total number of 216 country level of groups were defined. The deeper the colour, the higher the percentage of the isolates sourced from that country in continent level of groups. The background information of isolates is provided in Supplementary Table 1.

G35.C07 was the largest country specific group that contained 49 isolates from Colombia, followed by the G35.C05 (n=35) dominated in Colombia as well. These isolates from Colombia were mainly collected from the NCBI Bioproject PRJNA352848, which study contained the population structure of *H. pylori* in regional evolution in South America (Muñoz-Ramírez, et al. 2017). The isolates from group G35.C07 and G35.C05 were mainly found from Colombia, Mexico and Spain (**Figure 3**). This result provided the evidence that the *H. pylori* isolates were possibly transmitted from Spain and spread locally in South America and North America. In comparison, Australia and Switzerland were the largest countries of isolate source which isolates scattered across more than half of the country specific groups.

When comparing the percentage of isolates from different countries, those isolates from France, Germany, Malaysia, Nicaragua, Sweden, and UK were found scattered in more than one continent group, while isolates from Cambodia, Colombia, India, Peru, Spain and US were focused in one continent group when they were also found in other continent groups. More importantly, Australia and Switzerland were two countries that mostly found with scattered isolates in different regional specific groups.

Three clusters were observed in the percentage of different isolate sources at continent scale (G32 to G25 with red branches in Figure 3), consisting of groups from Europe and mixed continents. Specifically, those isolates from mixed groups were mainly sourced from European and Oceania countries, making this cluster as Europe-Oceania dominated. The second cluster was the mixed by Asian, Oceanian, European and mixed groups (G4 to G2 with green branches in Figure 3) but dominated by isolates from Australia and Asian countries. Therefore, cluster two was specified as Asian-Pacific cluster. The third cluster was formed by North American groups (G31 to G37 with purple branches in Figure 3), while South American branches was next to the North America cluster.

### Comparing with seven-gene MLST

Seven-gene MLST was implied to get sequence types (STs) of all 1,211 isolates. Unfortunately, due to the high mutation rate of the *H. pylori* strains, most of the seven-gene allele were only found with high similarity instead of an accurate type, as the result, a large number of isolates (n=876, 72.3%) were untyped in our dataset (**Supplementary Table 1 & Figure 1**). However, despite the most of undefined isolates, the typed isolates with exact ST number would still be grouped closely by minimum spanning tree.

### A user-friendly typing website

In order to support our *H. pylori* geographic typing tool, a user-friendly typing website was established and available at https://db.cngb.org/HPTT/. Our HTTP approach is compatible with any whole-genome sequencing (WGS) data with metadata (**Figure 4**). For the sequencing data from pure-cultured isolates, the assembled genomes can be directly submitted to our website. However, it is worth noting that sequences or assembled genomes needed to be extracted from metagenome samples before submission (Parks, et al. 2017; Olekhnovich, et al. 2019). Except for the sequenced genome data, the available assembled contigs from NCBI Sequence Read Archive (SRA) or assembly database (RefSeq), or other genome databases (e.g., European Nucleotide Achieve) can also be directly uploaded to our website. By using MUMmer alignment and blast process, the uploaded genome can be located to the closest matching genomes, further facilitating the possible transmission route analysis across the globe. In addition, our database can be also linked to the NCBI genome database, helping the user easily locate the metadata information from the available database (**Supplementary materials**).

**Figure 4.**
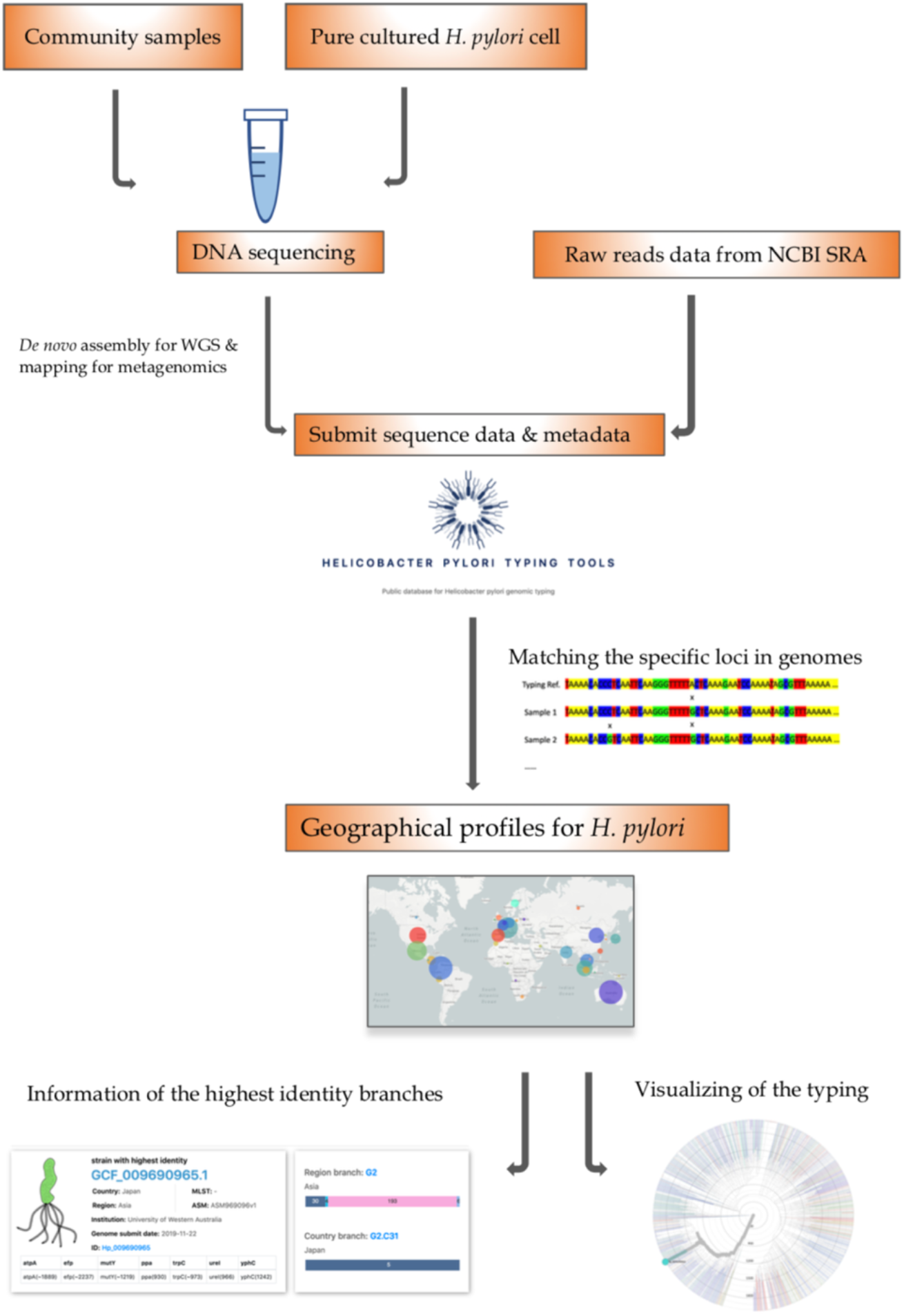
The HPTT workflow. The SNP based genotyping approach can be used with the Whole Genome Sequencing (WGS) data, which can be acquired in following ways: DNA can be extracted from a pure cultured bacterial cell with WGS data or a community sample with metagenomic sequencing data. After being sequenced by an appropriate platform, the assembled genomes can be directly submitted to our database. In addition, the public assembled data also can be directly submitted to our database. The downstream analyses of the aligned sequence data can be linked to the phylogenetic and geographic page.

Except for the typing tool, the Nextstrain framework was also embedded in our website. By clicking the uploaded genome number, information can be linked to the phylogenetic tree with corresponding continent and country. Possible evolution relationships and interactive located functions have made our typing tools easy to be applied and understood.

Ten genomes that newly uploaded to NCBI were downloaded and tested for the accuracy of the typing method and the efficiency of our website (**Supplementary Table 2**). Since our typing tool was established based on the MPS (Massive Parallel Sequencing) data, the first genome (GCF_002206465.1) sequenced by Pacbio was failed to be assigned groups. The rest nine genomes were typed successfully.

## Discussion

The epidemiological patterns of *H. pylori* isolates have been reported with specific geographic characteristics. In this study, the new typing webtool HPTT not only illustrated the population structure of *H. pylori* but also made the genomic typing easy to approach. In the continent level of typing, 1,112 isolates were grouped into 37 continent specific patterns. Except for 12 continent mixed groups, the rest could be defined as continent specific groups across the five continents. Isolates from Europe and Oceania were universally found in most of the continent-level groups (Europe 33/37, 89.19% and Oceania 26/37, 70.27%), illustrating that isolates from these two continents were widely spread across the world.

In the country level of typing, 1,045 isolates were grouped into 216 country level of groups. Most of the isolates were defined as country specific groups (168/216, 77.77%), while the rest of the isolates were grouped as country mixed groups (48/216, 22.22%). Australian and Swiss isolates were found to be widespread around the world, while isolates from Columbia was more regional specific. It has been reported that *H. pylori* in South America was originally transmitted from Spain (Muñoz-Ramírez, et al. 2017), this data perfectly aligned with our results in G35.C05 and G35.C07, giving the support of the accuracy of our genomic typing method.

In this study, except for the novel typing tool, a user-friendly website was also established. By using this typing tool, users can achieve fast and precise genomic typing, easily locating the possible origins and transmission events across the world. When located in the actual geographic group, it is easily for users to check the details of the corresponding composition of the branches in our database. The genome with the highest identity can be easily linked to the NCBI database as well as the visualization tool where the dynamic evolution of *H. pylori* was shown. At the same time, seven-gene MLST results were displayed for each genome in database.

The most interesting part of HPTT tool and methodology allow us to perform genome typing with assembled genomes from the metagenomics samples, as illustrated in Figure 4. Due to rapid mutation of *H. pylori*, it is most likely that the sample from one’s gut are heterogeneity in nature. The whole genome sequencing by combining sequencing libraries labelled with different barcodes on a meta sample, and a cultured pure isolate could yield enough data from one single run to perform the epidemiological surveillance of *H. pylori* on a global level to find the origins in evolution profile. An open-source assay protocol will be developed and shared in the future to combine with this HTTP tool to enable the epidemiological surveillance of *H. pylori*.

Although our typing tool filled the gap of genetic epidemiological surveillance of *H. pylori*, some of the functions still need to be improved. For example, cytotoxin-associated gene A (*cagA*) and *vacA* were the two crucial genes that reported to be correlated with geographic patterns of *H. pylori* (Yamaoka 2009; Breurec, et al. 2011). The *cagA* gene is one of the most important virulence genes in *H. pylori*, located at the end of cag pathogenicity island (cag PAI) that encodes 120–145 kDa CagA protein (Šterbenc, et al. 2019). Another virulence factor was vacuolating cytotoxin encoded by the gene *vacA* (Šterbenc, et al. 2019). The variation of these two genes were widely reported by the *H. pylori* groups that can reflect the genomic different for different geographic patterns. However, such rapid typing method on a website for these two genes are still lacking, which could be considered in the further HPTT version 2.

*H. pylori* is normally treated by the antibiotics without antimicrobial susceptibility testing (Pohl, et al. 2019). Antibiotics-resistant *H. pylori* has been reported related to several mutations within the genes *pbp1A*, *23S rRNA*, *gyrA*, *rdxA*, *frxA*, and *rpoB* (Domanovich-Asor, et al. 2021). In version 2, these antibiotics-resistant genes will be included in our second version despite an antibiotic-resistant specific tool was available now (Yusibova, et al. 2020). As more or more strains or isolates are deposited into our database with the geographic information, the HPTT tool will be evolute into a more powerful tool to associate the genomic typing information with its origin and phenotypes.

In summary, this work illustrates the efforts in global epidemiological study of *H. pylori* isolates. Two functions were designed for the web typing tool, one for genomic typing and the other for phylogenetic and geographic visualization. The accuracy of our genomic typing system was proved by ten unused genomes as well as another published study (Muñoz-Ramírez, et al. 2017). Together with the visualization tool, the genomic population structure of *H. pylori* with geographic documents were described. Future studies based on this approach will be expanded by the crucial virulence gene and antibiotic related genes. This tool would be beneficial for the surveillance of *H. pylori* for public health and the monitoring of its epidemic development.

## Acknowledgements

This study was supported by the Science, Technology and Innovation Commission of Shenzhen Municipality under grant (No. JCYJ20170412153155228). We thank China National GeneBank at Shenzhen for supporting this study.

## Data Availability

All assembled *H. pylori* genomes were downloaded from NCBI assembly database.

## Notes

### Competing Interest Statement

The authors have declared no competing interest.

