## Supplementary figures and images for "Whole-genome-based *Helicobacter pylori* geographic surveillance: a visualized and expandable webtool"

### Supplementary_Fig_1.pdf

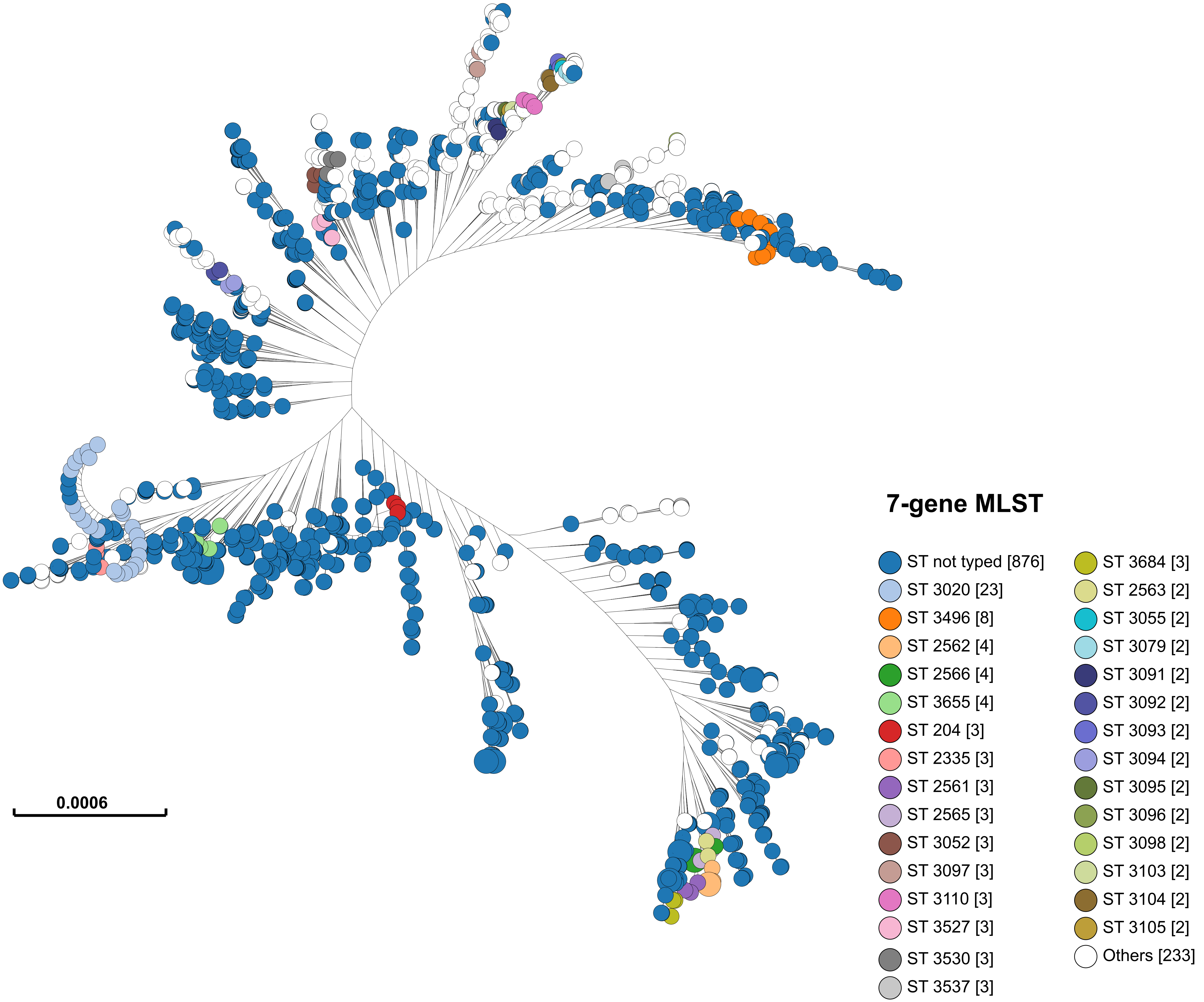
