## Supplementary_materials for "Whole-genome-based *Helicobacter pylori* geographic surveillance: a visualized and expandable webtool"

Supplementary Table 1. Strains information and typing results

| No._Of_Strains | Continent | Country | Ggroup | Cgroup | MLST types |
|---|---|---|---|---|---|
| Hp_002357575 | Asia | Japan | G2 | G2.C34 | - |
| Hp_003639245 | Europe | Spain | G25 | G25.C06 | - |
| Hp_002012505 | South America | Colombia | G35 | G35.C07 | 3103 |
| Hp_000285895 | Europe | France | G17 | G17.C01 | 3655 |
| Hp_003639785 | Europe | Spain | G32 | G32.C01 | - |
| Hp_001542505 | Asia | Malaysia | G2 | G2.C40 | - |
| Hp_003635825 | Oceania | Australia | G25 | G25.C05 | - |
| Hp_002533065 | South America | Colombia | G26 | G26.C04 | 3143 |
| Hp_000685705 | Oceania | Australia | G20 | NA | - |
| Hp_000499345 | Europe | Russia | G2 | G2.C01 | - |
| Hp_003637925 | Oceania | Australia | G25 | NA | - |
| Hp_004012975 | Europe | Switzerland | NA | 0.C12 | - |
| Hp_009737175 | North America | United States of America | G20 | G20.C03 | - |
| Hp_001275165 | Asia | Malaysia | G2 | G2.C24 | - |
| Hp_009741105 | North America | United States of America | G37 | NA | - |
| Hp_002092915 | Africa | Morocco | G37 | G37.C25 | 3496 |
| Hp_006439905 | Asia | Cambodia | NA | 0.C02 | - |
| Hp_003638105 | Oceania | Australia | G20 | G20.C07 | - |
| Hp_011141015 | North America | Mexico | G37 | G37.C11 | - |
| Hp_003638625 | Oceania | Australia | G20 | G20.C04 | 1144 |
| Hp_009690305 | Asia | China | G2 | G2.C04 | - |
| Hp_002012395 | South America | Colombia | G35 | G35.C07 | 3104 |
| Hp_011140875 | North America | Mexico | NA | NA | - |
| Hp_003637485 | Oceania | Australia | G28 | NA | - |
| Hp_003636005 | Oceania | Australia | G2 | G2.C09 | - |
| Hp_009691005 | Asia | China | G2 | G2.C20 | - |
| Hp_003637345 | Oceania | Australia | G2 | G2.C06 | - |
| Hp_003638285 | Oceania | Australia | G19 | G19.C03 | 1445 |
| Hp_003939085 | Europe | Belarus | G21 | NA | - |
| Hp_003060065 | Europe | France | NA | 0.C08 | - |
| Hp_000401335 | Asia | Malaysia | G3 | G3.C03 | - |
| Hp_002081955 | Africa | Morocco | G37 | G37.C25 | 3496 |
| Hp_003058635 | Europe | Sweden | G17 | G17.C02 | - |
| Hp_002012715 | South America | Colombia | G35 | G35.C07 | 3108 |
| Hp_000685625 | North America | United States of America | G5 | G5.C02 | - |
| Hp_004013875 | Europe | Switzerland | G25 | NA | - |
| Hp_002012935 | South America | Colombia | G26 | G26.C04 | 3129 |
| Hp_004012215 | Europe | Switzerland | G9 | NA | - |
| Hp_011140795 | North America | Mexico | NA | 0.C06 | - |
| Hp_002832045 | Asia | India | G8 | G8.C02 | - |
| Hp_003637165 | Oceania | Australia | G16 | G16.C04 | - |
| Hp_002952175 | North America | United States of America | G22 | G22.C04 | - |
| Hp_006440345 | Asia | Cambodia | G3 | G3.C02 | - |
| Hp_009690255 | Asia | China | G2 | G2.C19 | - |
| Hp_000591295 | Asia | China | G2 | G2.C42 | 2565 |
| Hp_002832195 | Asia | India | G11 | G11.C03 | - |
| Hp_002952235 | North America | United States of America | G22 | G22.C04 | 3020 |
| Hp_004013315 | Europe | Switzerland | G13 | G13.C02 | - |
| Hp_009690015 | Asia | China | G2 | G2.C30 | - |
| Hp_000685745 | Oceania | Australia | G13 | G13.C01 | - |
| Hp_002357655 | Asia | Japan | G2 | G2.C34 | - |
| Hp_004014535 | Europe | Switzerland | G18 | G18.C02 | - |
| Hp_009741095 | North America | United States of America | G31 | G31.C03 | - |
| Hp_002026585 | North America | Mexico | G34 | G34.C03 | 3520 |
| Hp_009690345 | Asia | China | G2 | G2.C36 | - |
| Hp_004013635 | Europe | Switzerland | G10 | G10.C01 | - |
| Hp_009689925 | Asia | China | G2 | G2.C32 | - |
| Hp_004088155 | Asia | China | G2 | G2.C07 | - |
| Hp_004013195 | Europe | Switzerland | G23 | G23.C02 | - |
| Hp_009741145 | North America | United States of America | G2 | G2.C49 | - |
| Hp_001185035 | Asia | Malaysia | G3 | NA | - |
| Hp_006339065 | Europe | Germany | G21 | NA | - |
| Hp_009689765 | Asia | China | G2 | G2.C10 | - |
| Hp_000689095 | North America | United States of America | G5 | G5.C02 | - |
| Hp_003635785 | Oceania | Australia | G37 | G37.C18 | - |
| Hp_006438425 | Asia | Cambodia | G2 | NA | - |
| Hp_003637305 | Oceania | Australia | G2 | NA | - |
| Hp_000448505 | South America | Colombia | G35 | G35.C06 | - |
| Hp_003059085 | Europe | Sweden | G21 | NA | - |
| Hp_003639855 | Europe | Spain | NA | 0.C07 | - |
| Hp_003635665 | Oceania | Australia | G37 | G37.C33 | - |
| Hp_000470075 | Europe | Germany | G14 | G14.C02 | - |
| Hp_003382505 | Europe | Spain | G23 | G23.C04 | - |
| Hp_003635945 | Oceania | Australia | G2 | NA | - |
| Hp_901687255 | Oceania | Australia | G9 | G9.C04 | - |
| Hp_003059615 | Europe | France | G37 | G37.C32 | - |
| Hp_006339235 | Europe | Germany | G25 | G25.C11 | - |
| Hp_009689685 | Asia | China | G2 | G2.C37 | - |
| Hp_011175325 | North America | Nicaragua | G37 | G37.C07 | - |
| Hp_006440225 | Asia | Cambodia | G2 | G2.C02 | - |
| Hp_003059525 | Europe | France | G28 | G28.C01 | - |
| Hp_006439785 | Asia | Cambodia | G2 | G2.C12 | - |
| Hp_002532995 | South America | Colombia | G26 | G26.C02 | 3081 |
| Hp_006438455 | Asia | Cambodia | G2 | G2.C03 | - |
| Hp_002357455 | Asia | Japan | G2 | G2.C34 | - |
| Hp_002832435 | Asia | India | G3 | G3.C12 | - |
| Hp_001446395 | Asia | Malaysia | G2 | G2.C14 | - |
| Hp_002357515 | Asia | Japan | G2 | G2.C34 | - |
| Hp_000401295 | Asia | India | G3 | NA | - |
| Hp_001940085 | Asia | Singapore | G2 | G2.C46 | - |
| Hp_000982695 | North America | United States of America | G37 | G37.C25 | 3496 |
| Hp_004013435 | Europe | Switzerland | G6 | NA | - |
| Hp_006438515 | Asia | Cambodia | G24 | G24.C01 | - |
| Hp_009689805 | Asia | China | G2 | G2.C36 | - |
| Hp_003639905 | Europe | Spain | G35 | G35.C04 | - |
| Hp_000444305 | Asia | Malaysia | G2 | G2.C41 | 3688 |
| Hp_000274965 | North America | United States of America | G37 | G37.C31 | - |
| Hp_001653395 | Asia | North Korea | G2 | G2.C17 | 45 |
| Hp_003059335 | Europe | France | NA | NA | - |
| Hp_001051495 | Asia | Malaysia | G11 | G11.C02 | - |

| No._Of_Strains | Continent | Country | Ggroup | Cgroup | MLST types |
|---|---|---|---|---|---|
| Hp_003636925 | Oceania | Australia | G37 | G37.C16 | - |
| Hp_004013765 | Europe | Switzerland | G27 | G27.C02 | - |
| Hp_006440195 | Asia | Cambodia | G2 | G2.C08 | - |
| Hp_006439735 | Asia | Cambodia | G3 | NA | - |
| Hp_002832345 | Asia | India | G11 | NA | - |
| Hp_006439915 | Asia | Cambodia | G3 | G3.C02 | - |
| Hp_002832485 | Asia | India | G3 | NA | - |
| Hp_003638115 | Oceania | Australia | G16 | G16.C03 | - |
| Hp_002026785 | North America | Mexico | G30 | G30.C01 | 3543 |
| Hp_003639495 | Europe | Spain | G23 | G23.C02 | - |
| Hp_006440045 | Asia | Cambodia | G4 | G4.C02 | - |
| Hp_004012465 | Europe | Switzerland | G25 | G25.C10 | - |
| Hp_003635585 | Oceania | Australia | G8 | G8.C02 | 625 |
| Hp_009689635 | Asia | China | G2 | G2.C41 | - |
| Hp_003059265 | Europe | France | NA | 0.C10 | - |
| Hp_000689115 | North America | United States of America | G5 | G5.C02 | - |
| Hp_004013925 | Europe | Switzerland | G37 | G37.C21 | - |
| Hp_003815525 | South America | Colombia | G35 | G35.C07 | 3054 |
| Hp_006337265 | Europe | Germany | G20 | G20.C02 | - |
| Hp_002831985 | Asia | India | G11 | NA | - |
| Hp_004012355 | Europe | Switzerland | G23 | G23.C01 | - |
| Hp_003636765 | Oceania | Australia | NA | NA | - |
| Hp_004014025 | Europe | Switzerland | G30 | NA | - |
| Hp_002026825 | North America | Mexico | G36 | G36.C02 | 3551 |
| Hp_002952575 | North America | United States of America | G22 | G22.C04 | 3020 |
| Hp_003637565 | Oceania | Australia | G16 | G16.C05 | - |
| Hp_002533105 | South America | Colombia | G35 | G35.C05 | 3144 |
| Hp_003639215 | Europe | Spain | NA | 0.C13 | - |
| Hp_003637635 | Oceania | Australia | G25 | NA | - |
| Hp_003815475 | South America | Colombia | G35 | G35.C05 | 3060 |
| Hp_002832595 | Asia | India | G3 | G3.C07 | - |
| Hp_003059405 | Europe | France | G17 | G17.C02 | - |
| Hp_002357775 | Asia | Japan | G2 | G2.C31 | - |
| Hp_003637045 | Oceania | Australia | G2 | G2.C01 | - |
| Hp_006337405 | Europe | Germany | G19 | G19.C01 | - |
| Hp_009740625 | North America | United States of America | NA | 0.C07 | - |
| Hp_008026815 | North America | Mexico | G37 | G37.C01 | 3521 |
| Hp_003058595 | Europe | Sweden | G22 | G22.C01 | - |
| Hp_004013305 | Europe | Switzerland | G13 | G13.C02 | - |
| Hp_002533285 | South America | Colombia | NA | 0.C20 | 3130 |
| Hp_002534615 | South America | Colombia | G37 | G37.C19 | 3141 |
| Hp_000296015 | Asia | Malaysia | G3 | G3.C08 | 3670 |
| Hp_011140925 | North America | Mexico | G36 | G36.C02 | 3538 |
| Hp_002533515 | South America | Colombia | G35 | G35.C06 | 3116 |
| Hp_008026375 | North America | Mexico | G31 | G31.C05 | - |
| Hp_006440005 | Asia | Cambodia | G4 | G4.C02 | - |
| Hp_011141045 | North America | Mexico | NA | 0.C07 | - |
| Hp_007794935 | Europe | United Kingdom | G22 | G22.C04 | 3020 |
| Hp_003635975 | Oceania | Australia | G2 | G2.C06 | - |
| Hp_002533455 | South America | Colombia | G35 | G35.C05 | 3120 |
| Hp_003636165 | Oceania | Australia | G29 | G29.C05 | - |
| Hp_003637875 | Oceania | Australia | G29 | G29.C05 | - |
| Hp_002533365 | South America | Colombia | G35 | G35.C05 | 3126 |
| Hp_001248025 | Asia | Malaysia | G3 | G3.C08 | - |
| Hp_009740365 | North America | United States of America | G37 | G37.C20 | - |
| Hp_009741065 | North America | United States of America | G31 | G31.C03 | - |
| Hp_000824865 | Europe | United Kingdom | G19 | G19.C02 | 594 |
| Hp_004014275 | Europe | Switzerland | G10 | G10.C02 | - |
| Hp_011140485 | North America | Mexico | G36 | G36.C03 | - |
| Hp_004013595 | Europe | Switzerland | G7 | G7.C03 | - |
| Hp_009690375 | Asia | China | G2 | NA | - |
| Hp_004013075 | Europe | Switzerland | G9 | G9.C05 | - |
| Hp_003637435 | Oceania | Australia | G2 | G2.C19 | - |
| Hp_003058845 | Europe | Belgium | G15 | G15.C01 | - |
| Hp_003059125 | Europe | United Kingdom | G16 | G16.C04 | - |
| Hp_011140805 | North America | Mexico | G34 | NA | - |
| Hp_003638235 | Oceania | Australia | G16 | G16.C02 | 1450 |
| Hp_002357395 | Asia | Japan | G2 | NA | - |
| Hp_009741175 | North America | United States of America | G2 | G2.C49 | - |
| Hp_003382515 | Europe | Spain | G23 | G23.C04 | - |
| Hp_003638175 | Oceania | Australia | G37 | G37.C18 | - |
| Hp_003059605 | Europe | France | NA | 0.C10 | - |
| Hp_004012855 | Europe | Switzerland | G23 | G23.C02 | - |
| Hp_002533425 | South America | Colombia | G35 | G35.C07 | 3122 |
| Hp_000444195 | Asia | China | G2 | G2.C42 | 2566 |
| Hp_000392515 | Asia | Malaysia | G11 | G11.C05 | - |
| Hp_002532725 | South America | Colombia | G35 | G35.C06 | 3101 |
| Hp_008681245 | North America | Mexico | G31 | NA | - |
| Hp_000315955 | North America | Canada | G1 | G1.C02 | 2935 |
| Hp_000392455 | Asia | Malaysia | G2 | NA | - |
| Hp_002533655 | South America | Colombia | G35 | G35.C07 | 3110 |
| Hp_001278525 | Asia | Kuwait | G24 | G24.C05 | - |
| Hp_009740655 | North America | United States of America | G31 | G31.C04 | - |
| Hp_000448425 | South America | Colombia | G37 | G37.C23 | - |
| Hp_010917675 | South America | Colombia | G37 | G37.C04 | - |
| Hp_002534555 | South America | Colombia | G35 | G35.C05 | - |
| Hp_002533985 | South America | Colombia | G35 | G35.C05 | 3078 |
| Hp_000688915 | North America | United States of America | G5 | G5.C02 | - |
| Hp_001653455 | South America | Angola | G28 | NA | 1909 |
| Hp_002012575 | South America | Colombia | G37 | G37.C08 | 3134 |
| Hp_004013845 | Europe | Switzerland | G9 | G9.C05 | - |
| Hp_008681355 | North America | Mexico | G33 | G33.C02 | - |
| Hp_009740985 | North America | United States of America | G37 | G37.C28 | - |
| Hp_011175275 | North America | Nicaragua | G37 | G37.C17 | - |
| Hp_004014495 | Europe | Switzerland | NA | NA | - |
| Hp_002832075 | Asia | India | G3 | G3.C11 | - |
| Hp_003636845 | Oceania | Australia | G15 | G15.C02 | - |
| Hp_011140955 | North America | Mexico | G25 | NA | - |
| Hp_003639735 | Europe | Spain | G31 | G31.C02 | - |
| Hp_009689815 | Asia | China | G2 | G2.C37 | - |

| | | | | | |
|---|---|---|---|---|---|
| Hp_002533775 | South America | Colombia | G35 | G35.C07 | 3100 |
| Hp_004012915 | Europe | Switzerland | G7 | G7.C01 | - |
| Hp_000802625 | Europe | Portugal | G27 | G27.C02 | - |
| Hp_004012735 | Europe | Switzerland | G25 | G25.C10 | - |
| Hp_000148875 | South America | Peru | G1 | G1.C01 | 3041 |
| Hp_006438625 | Asia | Cambodia | G3 | G3.C01 | - |
| Hp_009690225 | Asia | China | G2 | G2.C07 | - |
| Hp_004014185 | Europe | Switzerland | NA | 0.C07 | - |
| Hp_002952395 | North America | United States of America | G22 | G22.C03 | 3020 |
| Hp_002532615 | South America | Colombia | G26 | G26.C04 | 3129 |
| Hp_002533895 | South America | Colombia | G35 | G35.C07 | 3093 |
| Hp_011140435 | North America | Mexico | G37 | NA | - |
| Hp_003635905 | Oceania | Australia | NA | 0.C10 | - |
| Hp_003382545 | Europe | Spain | G23 | G23.C04 | - |
| Hp_000591335 | Asia | China | G2 | G2.C42 | 2565 |
| Hp_004323355 | Asia | Malaysia | G2 | G2.C38 | - |
| Hp_003637805 | Oceania | Australia | G9 | NA | - |
| Hp_000287835 | Asia | China | G2 | G2.C19 | 3673 |
| Hp_004014205 | Europe | Switzerland | G10 | G10.C03 | - |
| Hp_000817025 | Asia | China(Taiwan) | G2 | NA | - |
| Hp_002534385 | South America | Colombia | G37 | G37.C02 | 3090 |
| Hp_002534445 | South America | Colombia | G35 | G35.C07 | 3079 |
| Hp_003639325 | Europe | Spain | G32 | G32.C02 | - |
| Hp_008681215 | North America | Mexico | G36 | G36.C04 | - |
| Hp_004012135 | Europe | Switzerland | NA | 0.C11 | - |
| Hp_004012695 | Europe | Switzerland | G21 | G21.C01 | - |
| Hp_002533235 | South America | Colombia | G37 | G37.C08 | 3134 |
| Hp_003637225 | Oceania | Australia | G37 | G37.C18 | - |
| Hp_010917155 | South America | Colombia | G37 | G37.C08 | - |
| Hp_002533175 | South America | Colombia | G35 | G35.C05 | 3136 |
| Hp_003638425 | Oceania | Australia | G12 | G12.C02 | - |
| Hp_001189755 | Asia | Malaysia | G2 | G2.C38 | - |
| Hp_003637685 | Oceania | Australia | G29 | G29.C05 | - |
| Hp_009690075 | Asia | China | G2 | G2.C20 | - |
| Hp_003639665 | Europe | Spain | G37 | G37.C20 | - |
| Hp_003635855 | Oceania | Australia | G16 | G16.C02 | - |
| Hp_000444345 | Asia | Malaysia | G2 | G2.C27 | 3680 |
| Hp_001433495 | North America | Mexico | G27 | G27.C02 | - |
| Hp_002534535 | South America | Colombia | G26 | G26.C03 | - |
| Hp_003636395 | Oceania | Australia | G29 | G29.C04 | - |
| Hp_002026835 | North America | Mexico | G36 | G36.C02 | 3519 |
| Hp_002832505 | Asia | India | G7 | G7.C04 | - |
| Hp_001653435 | Oceania | Australia | NA | NA | 1047 |
| Hp_003637625 | Oceania | Australia | G28 | NA | - |
| Hp_002012795 | South America | Colombia | G35 | G35.C05 | 3120 |
| Hp_008681335 | North America | Mexico | G35 | G35.C06 | - |
| Hp_003637105 | Oceania | Australia | G24 | G24.C07 | - |
| Hp_009740565 | North America | United States of America | G37 | G37.C29 | - |
| Hp_002533195 | South America | Colombia | NA | 0.C07 | 3135 |
| Hp_000740115 | Asia | China | G2 | G2.C04 | - |
| Hp_000451005 | Asia | Malaysia | G11 | G11.C05 | - |
| Hp_011140715 | North America | Mexico | G31 | G31.C01 | - |
| Hp_003637515 | Oceania | Australia | G2 | G2.C08 | - |
| Hp_009690145 | Asia | China | G2 | G2.C21 | - |
| Hp_001538305 | Asia | Malaysia | G2 | G2.C18 | - |
| Hp_008728675 | North America | Mexico | G30 | G30.C01 | - |
| Hp_000591215 | Asia | China | G2 | G2.C43 | 2563 |
| Hp_008027235 | North America | Mexico | G34 | G34.C01 | - |
| Hp_001446435 | Asia | Malaysia | G2 | G2.C18 | - |
| Hp_003639265 | Europe | Spain | G25 | G25.C06 | - |
| Hp_004013395 | Europe | Switzerland | G27 | G27.C01 | - |
| Hp_003637285 | Oceania | Australia | G20 | G20.C05 | - |
| Hp_002832115 | Asia | India | G3 | G3.C12 | - |
| Hp_003638485 | Oceania | Australia | G24 | G24.C06 | - |
| Hp_003637745 | Oceania | Australia | G20 | G20.C07 | - |
| Hp_003638345 | Oceania | Australia | G37 | G37.C28 | - |
| Hp_011140935 | North America | Mexico | G36 | G36.C04 | - |
| Hp_011140545 | North America | Mexico | G36 | G36.C03 | - |
| Hp_008681525 | North America | Mexico | G30 | G30.C01 | - |
| Hp_000281675 | Asia | Malaysia | G3 | NA | - |
| Hp_006440145 | Asia | Cambodia | NA | 0.C01 | - |
| Hp_002026675 | North America | Mexico | G31 | G31.C01 | 3536 |
| Hp_002952375 | North America | United States of America | G22 | G22.C04 | 3020 |
| Hp_004117955 | Asia | China(Taiwan) | G2 | NA | - |
| Hp_002534325 | South America | Colombia | G26 | G26.C04 | 3111 |
| Hp_003637365 | Oceania | Australia | G19 | G19.C03 | - |
| Hp_004012835 | Europe | Switzerland | G27 | G27.C02 | - |
| Hp_003639445 | Europe | Spain | NA | 0.C13 | - |
| Hp_006440095 | Asia | Cambodia | G3 | G3.C01 | - |
| Hp_003639385 | Europe | Spain | G25 | G25.C08 | - |
| Hp_002832395 | Asia | India | G3 | G3.C13 | - |
| Hp_009689705 | Asia | China | G2 | G2.C09 | - |
| Hp_003636505 | Oceania | Australia | G2 | NA | - |
| Hp_004013115 | Europe | Switzerland | G2 | G2.C34 | - |
| Hp_001248035 | Asia | Malaysia | G3 | G3.C05 | - |
| Hp_000259235 | Europe | Spain | G26 | G26.C02 | 3021 |
| Hp_003855475 | Asia | China | G2 | G2.C28 | - |
| Hp_000689015 | North America | United States of America | G5 | G5.C02 | 204 |
| Hp_005796115 | Europe | Poland | G17 | G17.C01 | 3655 |
| Hp_004013505 | Europe | Switzerland | G13 | G13.C02 | - |
| Hp_008033015 | North America | Mexico | G30 | G30.C01 | - |
| Hp_009740315 | North America | United States of America | G37 | G37.C28 | - |
| Hp_004014605 | Europe | Switzerland | G12 | NA | - |
| Hp_001446375 | Asia | Malaysia | G2 | G2.C18 | - |
| Hp_001998045 | South America | Colombia | G37 | G37.C15 | 3508 |
| Hp_008027775 | North America | Mexico | G31 | G31.C05 | - |
| Hp_001939635 | Asia | Singapore | G3 | G3.C09 | - |
| Hp_002832545 | Asia | India | G8 | G8.C01 | - |
| Hp_000256035 | Asia | India | G13 | NA | - |
| Hp_000287735 | Asia | China | G2 | G2.C28 | 2164 |
| Hp_003638305 | Oceania | Australia | NA | NA | - |
| Hp_001653375 | Asia | India | G2 | G2.C01 | - |
| Hp_008681475 | North America | Mexico | G37 | G37.C12 | - |

| | | | | | |
|---|---|---|---|---|---|
| Hp_003639915 | Europe | Spain | G35 | NA | - |
| Hp_011175135 | North America | Nicaragua | G37 | G37.C10 | - |
| Hp_000277405 | South America | Peru | G1 | G1.C02 | 3044 |
| Hp_003640045 | Europe | Spain | NA | 0.C11 | - |
| Hp_001991095 | Oceania | Australia | G9 | G9.C04 | - |
| Hp_004012905 | Europe | Switzerland | G24 | G24.C05 | - |
| Hp_002027025 | North America | Mexico | G33 | G33.C02 | 3525 |
| Hp_008032615 | North America | Mexico | NA | 0.C14 | 3527 |
| Hp_003635915 | Oceania | Australia | G37 | G37.C24 | - |
| Hp_001189835 | Asia | Malaysia | G3 | NA | - |
| Hp_003059965 | Europe | France | G10 | G10.C02 | - |
| Hp_004013225 | Europe | Switzerland | G20 | NA | - |
| Hp_011140845 | North America | Mexico | G31 | G31.C05 | - |
| Hp_002532605 | South America | Colombia | G35 | G35.C05 | 3137 |
| Hp_003635735 | Oceania | Australia | G29 | G29.C05 | - |
| Hp_003058805 | Europe | United Kingdom | G19 | NA | - |
| Hp_002026665 | North America | Mexico | G33 | G33.C02 | 3535 |
| Hp_004014585 | Europe | Switzerland | G24 | G24.C02 | - |
| Hp_003637375 | Oceania | Australia | G37 | G37.C33 | - |
| Hp_004012325 | Europe | Switzerland | G23 | G23.C03 | - |
| Hp_008032475 | North America | Mexico | G30 | G30.C01 | - |
| Hp_002952455 | North America | United States of America | G22 | G22.C03 | 3020 |
| Hp_009740865 | North America | United States of America | G34 | NA | - |
| Hp_002013035 | South America | Colombia | G35 | G35.C07 | 3095 |
| Hp_003636225 | Oceania | Australia | G16 | G16.C03 | 1763 |
| Hp_002012425 | South America | Colombia | G26 | G26.C04 | - |
| Hp_004013805 | Europe | Switzerland | G10 | G10.C03 | - |
| Hp_001189675 | Asia | Malaysia | G3 | G3.C03 | - |
| Hp_002952515 | North America | United States of America | G22 | G22.C04 | 3020 |
| Hp_000287755 | Asia | China | G2 | G2.C23 | 2128 |
| Hp_006440435 | Asia | Cambodia | G2 | NA | - |
| Hp_002357715 | Asia | Japan | G2 | G2.C31 | - |
| Hp_006438545 | Asia | Cambodia | G4 | G4.C02 | - |
| Hp_008032995 | North America | Mexico | G36 | G36.C01 | - |
| Hp_004012535 | Europe | Switzerland | G10 | G10.C03 | - |
| Hp_002012655 | South America | Colombia | G35 | G35.C07 | - |
| Hp_003058685 | Europe | Sweden | G20 | G20.C06 | - |
| Hp_004012665 | Europe | Switzerland | G10 | G10.C02 | - |
| Hp_000277385 | South America | Peru | G1 | G1.C02 | 3045 |
| Hp_002534295 | South America | Colombia | G37 | G37.C09 | 3056 |
| Hp_010917555 | South America | Colombia | G37 | G37.C09 | - |
| Hp_003636805 | Oceania | Australia | G2 | G2.C44 | - |
| Hp_009740575 | North America | United States of America | G37 | NA | - |
| Hp_002533575 | South America | Colombia | G34 | G34.C04 | 3113 |
| Hp_006338945 | Europe | Germany | G20 | G20.C02 | - |
| Hp_003637115 | Oceania | Australia | G29 | G29.C02 | - |
| Hp_002832285 | Asia | India | G3 | G3.C06 | - |
| Hp_001189695 | Asia | Malaysia | G2 | NA | - |
| Hp_002026445 | North America | Mexico | G36 | G36.C01 | 3518 |
| Hp_011175085 | North America | Nicaragua | G27 | G27.C02 | - |
| Hp_004014335 | Europe | Switzerland | NA | NA | - |
| Hp_003058625 | Europe | Sweden | G20 | G20.C01 | - |
| Hp_004013365 | Europe | Switzerland | G3 | G3.C14 | - |
| Hp_009690065 | Asia | China | G2 | G2.C20 | - |
| Hp_003639675 | Europe | Spain | G26 | G26.C03 | - |
| Hp_008032975 | North America | Mexico | G35 | G35.C03 | - |
| Hp_006337465 | Europe | Germany | G14 | NA | - |
| Hp_002534135 | South America | Colombia | G35 | G35.C07 | 3071 |
| Hp_004012755 | Europe | Switzerland | G25 | G25.C05 | - |
| Hp_000296095 | Asia | Malaysia | G3 | NA | - |
| Hp_003637505 | Oceania | Australia | G20 | G20.C07 | - |
| Hp_004012265 | Europe | Switzerland | G10 | G10.C02 | - |
| Hp_003058745 | Europe | Sweden | G22 | G22.C01 | - |
| Hp_003636455 | Oceania | Australia | G2 | G2.C35 | - |
| Hp_002533595 | South America | Colombia | G37 | G37.C03 | 3512 |
| Hp_002831955 | Asia | India | G5 | NA | - |
| Hp_001542485 | Asia | Malaysia | G3 | G3.C14 | - |
| Hp_004012445 | Europe | Switzerland | G25 | G25.C05 | - |
| Hp_008026175 | North America | Mexico | G35 | NA | - |
| Hp_011141115 | North America | Mexico | G37 | G37.C07 | 3537 |
| Hp_003636745 | Oceania | Australia | G2 | G2.C09 | - |
| Hp_004013625 | Europe | Switzerland | G17 | G17.C02 | - |
| Hp_003059245 | Europe | France | G29 | G29.C01 | - |
| Hp_003636285 | Oceania | Australia | G16 | G16.C03 | 1055 |
| Hp_003635555 | Oceania | Australia | G20 | G20.C04 | 1733 |
| Hp_002012485 | South America | Colombia | G35 | G35.C07 | 3110 |
| Hp_004323475 | Asia | Malaysia | G25 | G25.C07 | - |
| Hp_004013745 | Europe | Switzerland | G24 | G24.C05 | - |
| Hp_002357435 | Asia | Japan | G2 | G2.C34 | - |
| Hp_001185025 | Asia | Malaysia | G2 | G2.C47 | 3684 |
| Hp_002026755 | North America | Mexico | G35 | G35.C01 | 3541 |
| Hp_009740305 | North America | United States of America | G25 | NA | - |
| Hp_003060165 | Europe | France | G25 | NA | - |
| Hp_002203985 | Asia | China | G2 | G2.C44 | - |
| Hp_003636105 | Oceania | Australia | G4 | G4.C01 | - |
| Hp_004323525 | Oceania | Australia | G29 | G29.C05 | - |
| Hp_000740095 | Asia | China | G2 | G2.C04 | - |
| Hp_002533765 | South America | Colombia | G35 | G35.C07 | 3099 |
| Hp_004012155 | Europe | Switzerland | G17 | G17.C02 | - |
| Hp_011174985 | North America | Nicaragua | G24 | G24.C06 | - |
| Hp_002027085 | North America | Mexico | G33 | G33.C01 | 3547 |
| Hp_011140625 | North America | Mexico | G30 | G30.C01 | - |
| Hp_006440395 | Asia | Cambodia | G4 | G4.C02 | - |
| Hp_003058705 | Europe | Sweden | NA | 0.C05 | - |
| Hp_004014475 | Europe | Switzerland | G6 | NA | - |
| Hp_011175295 | North America | Nicaragua | G37 | NA | 3574 |
| Hp_000448485 | South America | Colombia | G35 | G35.C06 | - |
| Hp_011140745 | North America | Mexico | G35 | G35.C03 | - |
| Hp_003637935 | Oceania | Australia | G20 | G20.C04 | - |
| Hp_900450825 | Europe | Belgium | G25 | G25.C06 | - |
| Hp_003637545 | Oceania | Australia | G7 | G7.C02 | - |
| Hp_003637715 | Oceania | Australia | G25 | G25.C01 | - |
| Hp_002533925 | South America | Colombia | G26 | G26.C04 | 3088 |

| | | | | | |
|---|---|---|---|---|---|
| Hp_000296115 | Asia | Malaysia | G3 | G3.C05 | - |
| Hp_001940025 | Asia | Singapore | G2 | G2.C47 | - |
| Hp_004012555 | Europe | Switzerland | G14 | G14.C01 | - |
| Hp_000148895 | South America | Peru | G1 | G1.C02 | 3040 |
| Hp_003058715 | Europe | Sweden | G17 | G17.C02 | - |
| Hp_002012635 | South America | Colombia | G35 | G35.C05 | 3136 |
| Hp_009690105 | Asia | China | G2 | NA | - |
| Hp_002012565 | South America | Colombia | G35 | G35.C05 | 3117 |
| Hp_009689745 | Asia | China | G2 | G2.C29 | - |
| Hp_003636935 | Oceania | Australia | G28 | NA | - |
| Hp_003636545 | Oceania | Australia | G11 | G11.C01 | - |
| Hp_011141095 | North America | Mexico | G37 | G37.C01 | 3524 |
| Hp_003639405 | Europe | Spain | NA | 0.C06 | - |
| Hp_003636065 | Oceania | Australia | G2 | G2.C11 | - |
| Hp_006440105 | Asia | Cambodia | NA | NA | - |
| Hp_011141145 | North America | Mexico | G30 | G30.C01 | 3530 |
| Hp_006338415 | Europe | Germany | G19 | G19.C01 | - |
| Hp_004013065 | Europe | Switzerland | NA | 0.C10 | - |
| Hp_002952435 | North America | United States of America | G22 | G22.C03 | 3020 |
| Hp_002013055 | South America | Colombia | NA | 0.C19 | 3109 |
| Hp_002532975 | South America | Colombia | G35 | G35.C06 | 3106 |
| Hp_004013715 | Europe | Switzerland | G37 | G37.C24 | - |
| Hp_011105805 | North America | Mexico | G37 | G37.C11 | - |
| Hp_006439965 | Asia | Cambodia | G4 | G4.C02 | - |
| Hp_004013545 | Europe | Switzerland | G25 | NA | - |
| Hp_009689625 | Asia | China | G2 | G2.C40 | - |
| Hp_001939315 | Asia | Singapore | G2 | G2.C29 | - |
| Hp_009741165 | North America | United States of America | G37 | G37.C30 | - |
| Hp_002012875 | South America | Colombia | G37 | G37.C05 | 3124 |
| Hp_006439645 | Asia | Cambodia | G2 | G2.C12 | - |
| Hp_006440285 | Asia | Cambodia | G2 | G2.C03 | - |
| Hp_002026555 | North America | Mexico | G35 | G35.C02 | - |
| Hp_003638165 | Oceania | Australia | G20 | G20.C05 | - |
| Hp_003638595 | Oceania | Australia | G29 | G29.C04 | - |
| Hp_008027395 | North America | Mexico | G34 | G34.C01 | - |
| Hp_002533815 | South America | Colombia | G37 | G37.C08 | 3098 |
| Hp_003059025 | Europe | Sweden | G20 | G20.C06 | - |
| Hp_009740815 | North America | United States of America | G37 | G37.C31 | - |
| Hp_004013175 | Europe | Switzerland | G25 | G25.C08 | - |
| Hp_003635965 | Oceania | Australia | G10 | G10.C03 | - |
| Hp_003637865 | Oceania | Australia | G37 | G37.C18 | - |
| Hp_003635645 | Oceania | Australia | G37 | G37.C04 | - |
| Hp_009690425 | Asia | China | G2 | NA | - |
| Hp_000689075 | North America | United States of America | G5 | G5.C02 | - |
| Hp_008033075 | North America | Mexico | G33 | G33.C02 | 3535 |
| Hp_009689985 | Asia | China | NA | NA | - |
| Hp_002533375 | South America | Colombia | G37 | G37.C05 | 3124 |
| Hp_002534185 | South America | Colombia | G35 | G35.C07 | 3068 |
| Hp_003639605 | Europe | Spain | G32 | G32.C02 | - |
| Hp_006440425 | Asia | Cambodia | G2 | G2.C14 | - |
| Hp_002027125 | North America | Mexico | G31 | G31.C05 | 3548 |
| Hp_003637575 | Oceania | Australia | G2 | G2.C20 | - |
| Hp_002952195 | North America | United States of America | G22 | G22.C04 | - |
| Hp_009690985 | Asia | China | G2 | G2.C19 | - |
| Hp_003640145 | Europe | Spain | G36 | G36.C03 | - |
| Hp_011105765 | North America | Mexico | G36 | G36.C04 | 3546 |
| Hp_000591275 | Asia | China | G2 | G2.C42 | 2565 |
| Hp_002952635 | North America | United States of America | G22 | G22.C03 | 3020 |
| Hp_011175035 | North America | Nicaragua | G37 | G37.C15 | - |
| Hp_004014385 | Europe | Switzerland | G10 | G10.C03 | - |
| Hp_003640095 | Europe | Spain | NA | 0.C09 | - |
| Hp_002832175 | Asia | India | G3 | G3.C10 | - |
| Hp_002831875 | Asia | India | G11 | G11.C03 | - |
| Hp_000591135 | Asia | China | G2 | G2.C43 | 2562 |
| Hp_000521245 | Asia | China | G2 | G2.C19 | 2139 |
| Hp_004012475 | Europe | Switzerland | G25 | G25.C05 | - |
| Hp_000224575 | Asia | India | G3 | NA | 3031 |
| Hp_003638225 | Oceania | Australia | G25 | NA | - |
| Hp_011140895 | North America | Mexico | G35 | G35.C01 | - |
| Hp_002154535 | Africa | Morocco | G37 | G37.C25 | 3496 |
| Hp_002013135 | South America | Colombia | G35 | G35.C07 | - |
| Hp_003070765 | Europe | France | G27 | G27.C01 | - |
| Hp_000444745 | Asia | Malaysia | G2 | NA | 3685 |
| Hp_002532915 | South America | Colombia | G26 | G26.C02 | 3083 |
| Hp_004323385 | Oceania | Australia | G25 | G25.C04 | - |
| Hp_004323445 | Oceania | Australia | G10 | G10.C01 | - |
| Hp_002012815 | South America | Colombia | G35 | G35.C07 | 3121 |
| Hp_002532735 | South America | Colombia | G35 | G35.C07 | 3059 |
| Hp_001446315 | Asia | Malaysia | G2 | G2.C45 | - |
| Hp_002357595 | Asia | Japan | G2 | G2.C33 | - |
| Hp_003635885 | Oceania | Australia | G2 | G2.C39 | - |
| Hp_003637985 | Oceania | Australia | G2 | G2.C32 | - |
| Hp_000444365 | Asia | Malaysia | G2 | G2.C07 | 3681 |
| Hp_003639965 | Europe | Spain | G24 | NA | - |
| Hp_009689865 | Asia | China | G2 | NA | - |
| Hp_008281125 | North America | Mexico | G36 | G36.C04 | - |
| Hp_008681415 | North America | Mexico | G37 | G37.C12 | - |
| Hp_003639725 | Europe | Spain | NA | 0.C11 | - |
| Hp_002534245 | South America | Colombia | G35 | G35.C05 | 3058 |
| Hp_003640105 | Europe | Spain | G9 | G9.C05 | - |
| Hp_011174935 | North America | Nicaragua | G37 | G37.C10 | - |
| Hp_002533295 | South America | Colombia | NA | 0.C06 | 3128 |
| Hp_002012555 | South America | Colombia | G34 | G34.C04 | 3113 |
| Hp_001653475 | Oceania | Papua New Guinea | NA | NA | 1161 |
| Hp_003638405 | Oceania | Australia | G16 | G16.C05 | - |
| Hp_006338045 | Europe | Germany | G25 | G25.C08 | - |
| Hp_004012635 | Europe | Switzerland | G15 | G15.C02 | - |
| Hp_003058585 | Europe | Sweden | G20 | G20.C02 | - |
| Hp_004012565 | Europe | Switzerland | G14 | G14.C01 | - |
| Hp_003637205 | Oceania | Australia | G20 | G20.C07 | - |
| Hp_000392475 | Asia | Singapore | G2 | G2.C26 | - |
| Hp_002533675 | South America | Colombia | G35 | G35.C07 | 3110 |
| Hp_011175015 | North America | Nicaragua | G37 | G37.C17 | - |
| Hp_011140515 | North America | Mexico | NA | 0.C14 | 3527 |
| Hp_002831845 | Asia | India | G3 | NA | - |
| Hp_009740925 | North America | United States of America | G37 | G37.C27 | - |
| Hp_001998165 | South America | Colombia | G33 | G33.C02 | 3500 |
| Hp_001189735 | Asia | Malaysia | G6 | G6.C01 | - |
| Hp_002832145 | Asia | India | G4 | G4.C01 | - |
| Hp_003637065 | Oceania | Australia | G8 | G8.C01 | - |
| Hp_009741045 | North America | United States of America | G37 | G37.C14 | - |
| Hp_000591355 | Asia | China | G2 | G2.C42 | 2566 |
| Hp_002013185 | South America | Colombia | G35 | G35.C05 | 3517 |
| Hp_004323335 | Asia | Malaysia | G3 | G3.C04 | - |
| Hp_002357375 | Asia | Japan | G2 | G2.C38 | - |
| Hp_009798325 | Africa | Morocco | G37 | G37.C25 | 3496 |
| Hp_003638045 | Oceania | Australia | G29 | G29.C02 | - |
| Hp_003636705 | Oceania | Australia | G2 | G2.C18 | - |
| Hp_004012405 | Europe | Switzerland | G23 | G23.C02 | - |
| Hp_006439995 | Asia | Cambodia | NA | NA | - |
| Hp_000444175 | Asia | China | G2 | G2.C43 | 2561 |
| Hp_003638195 | Oceania | Australia | G20 | G20.C07 | - |
| Hp_011140365 | North America | Mexico | G26 | G26.C01 | 3532 |
| Hp_003638565 | Oceania | Australia | G21 | G21.C02 | 758 |
| Hp_003059185 | Europe | United Kingdom | G37 | G37.C22 | - |
| Hp_002013205 | South America | Colombia | NA | 0.C20 | 3130 |
| Hp_003636145 | Oceania | Australia | G2 | G2.C09 | 3259 |
| Hp_002357815 | Asia | Japan | G2 | G2.C17 | - |
| Hp_009740485 | North America | United States of America | G37 | G37.C16 | - |
| Hp_003636095 | Oceania | Australia | G9 | G9.C05 | - |
| Hp_001549855 | Asia | China(Taiwan) | G2 | G2.C49 | - |
| Hp_009740805 | North America | United States of America | G37 | NA | - |
| Hp_011140425 | North America | Mexico | G37 | G37.C06 | - |
| Hp_003957395 | Asia | South Korea | G2 | NA | - |
| Hp_004323415 | Oceania | Australia | G2 | G2.C03 | - |
| Hp_000401355 | Asia | Malaysia | G2 | G2.C07 | 3678 |
| Hp_004014675 | Europe | Switzerland | G9 | NA | - |
| Hp_003058865 | Europe | Belgium | G11 | NA | - |
| Hp_003639335 | Europe | Spain | G25 | G25.C05 | - |
| Hp_004014295 | Europe | Switzerland | G16 | G16.C05 | - |
| Hp_003059905 | Europe | France | G12 | G12.C02 | - |
| Hp_002952595 | North America | United States of America | G22 | G22.C03 | 3020 |
| Hp_008032735 | North America | Mexico | G37 | G37.C16 | - |
| Hp_003637175 | Oceania | Australia | G32 | G32.C01 | - |
| Hp_004012125 | Europe | Switzerland | G25 | NA | - |
| Hp_011105795 | North America | Mexico | G37 | G37.C11 | 3545 |
| Hp_001940185 | Asia | Singapore | G2 | G2.C20 | - |
| Hp_001189585 | Asia | Malaysia | G11 | NA | - |
| Hp_002533225 | South America | Colombia | G35 | G35.C07 | 3133 |
| Hp_011140655 | North America | Mexico | G37 | G37.C11 | - |
| Hp_003639615 | Europe | Spain | NA | 0.C06 | - |
| Hp_002357795 | Asia | Japan | G2 | G2.C39 | - |
| Hp_003637235 | Oceania | Australia | G20 | G20.C01 | - |
| Hp_002832575 | Asia | India | G7 | G7.C04 | - |
| Hp_004012605 | Europe | Switzerland | G17 | G17.C02 | - |
| Hp_001051445 | Asia | Malaysia | G2 | G2.C14 | - |
| Hp_002534585 | South America | Colombia | G35 | G35.C06 | 3053 |
| Hp_002026885 | North America | Mexico | G31 | G31.C01 | 3523 |
| Hp_002534055 | South America | Colombia | NA | 0.C16 | 3052 |
| Hp_002534215 | South America | Colombia | G35 | G35.C06 | 3065 |
| Hp_004013865 | Europe | Switzerland | G2 | G2.C09 | - |
| Hp_002533685 | South America | Colombia | G35 | G35.C07 | 3108 |
| Hp_000091345 | Europe | France | G9 | NA | 3022 |
| Hp_003639525 | Europe | Spain | G12 | G12.C01 | - |
| Hp_009740685 | North America | United States of America | G37 | G37.C30 | - |
| Hp_004013655 | Europe | Switzerland | G23 | G23.C02 | - |
| Hp_002357315 | Asia | Japan | G2 | G2.C31 | - |
| Hp_002026645 | North America | Mexico | G31 | G31.C05 | 3534 |
| Hp_009740845 | North America | United States of America | G37 | G37.C27 | - |
| Hp_002831925 | Asia | India | G15 | G15.C02 | - |
| Hp_002533845 | South America | Colombia | G35 | G35.C05 | 3097 |
| Hp_011175335 | North America | Nicaragua | G33 | G33.C01 | - |
| Hp_003059535 | Europe | France | G12 | NA | - |
| Hp_003636985 | Oceania | Australia | G29 | G29.C05 | - |
| Hp_002534495 | South America | Colombia | G35 | G35.C06 | 3076 |
| Hp_002012805 | South America | Colombia | G26 | G26.C04 | - |
| Hp_003635615 | Oceania | Australia | G11 | G11.C01 | - |
| Hp_002356595 | North America | United States of America | G2 | G2.C33 | - |
| Hp_002532905 | South America | Colombia | G35 | G35.C05 | 3080 |
| Hp_002026725 | North America | Mexico | G36 | G36.C02 | 3538 |
| Hp_900478295 | Oceania | Australia | G22 | G22.C02 | - |
| Hp_000295995 | Asia | Malaysia | G2 | G2.C09 | - |
| Hp_002012955 | South America | Colombia | G35 | G35.C06 | 3127 |
| Hp_003058985 | Europe | Sweden | NA | 0.C04 | - |
| Hp_002012685 | South America | Colombia | G35 | G35.C05 | 3138 |
| Hp_001446305 | Asia | Malaysia | G2 | G2.C49 | - |
| Hp_003058655 | Europe | Sweden | NA | 0.C04 | - |
| Hp_000688995 | North America | United States of America | G5 | G5.C02 | - |
| Hp_009737165 | North America | United States of America | G20 | G20.C03 | 2335 |
| Hp_002952255 | North America | United States of America | G22 | G22.C04 | 3020 |
| Hp_004014415 | Europe | Switzerland | G13 | G13.C02 | - |
| Hp_000824925 | Europe | United Kingdom | G19 | G19.C02 | 594 |
| Hp_006337325 | Europe | Germany | G24 | G24.C08 | - |
| Hp_000472305 | North America | Mexico | G31 | G31.C05 | - |
| Hp_002357635 | Asia | Japan | G2 | G2.C34 | - |
| Hp_003059325 | Europe | France | G19 | NA | - |
| Hp_003638375 | Oceania | Australia | G14 | G14.C02 | - |
| Hp_003637775 | Oceania | Australia | G29 | G29.C02 | - |
| Hp_003639995 | Europe | Spain | G32 | G32.C02 | - |
| Hp_006438565 | Asia | Cambodia | G4 | G4.C02 | - |
| Hp_002209285 | Europe | Switzerland | G9 | G9.C04 | - |
| Hp_009690235 | Asia | China | G2 | G2.C16 | - |
| Hp_004014375 | Europe | Switzerland | G9 | G9.C03 | - |
| Hp_003637055 | Oceania | Australia | G16 | G16.C05 | - |
| Hp_000448445 | South America | Colombia | G35 | G35.C06 | 3511 |
| Hp_003639565 | Europe | Spain | NA | NA | - |

| | | | | | | | | | | | |
|---|---|---|---|---|---|---|---|---|---|---|---|
| Hp_002952615 | North America | United States of America | G22 | G22.C04 | 3020 | Hp_003638415 | Oceania | Australia | G25 | G25.C08 | - |
| Hp_001940175 | Asia | Singapore | G2 | NA | - | Hp_003639195 | Europe | Spain | G27 | G27.C01 | - |
| Hp_001549715 | Asia | China(Taiwan) | G2 | NA | - | Hp_008032935 | North America | Mexico | G24 | G24.C06 | - |
| Hp_003711165 | Asia | Japan | G3 | NA | - | Hp_000766025 | Asia | China(Taiwan) | G2 | G2.C15 | 3677 |
| Hp_002832215 | Asia | India | G3 | G3.C10 | - | Hp_003639635 | Europe | Spain | G35 | G35.C04 | - |
| Hp_002832055 | Asia | India | G3 | G3.C10 | - | Hp_004013825 | Europe | Switzerland | G18 | G18.C02 | - |
| Hp_003639165 | Europe | Spain | G37 | NA | - | Hp_002534175 | South America | Colombia | G26 | G26.C04 | 3066 |
| Hp_006438645 | Asia | Cambodia | G4 | G4.C02 | - | Hp_003636825 | Oceania | Australia | G30 | NA | - |
| Hp_003637825 | Oceania | Australia | G37 | G37.C22 | - | Hp_000148855 | South America | Peru | NA | NA | 3050 |
| Hp_004013215 | Europe | Switzerland | G17 | G17.C02 | - | Hp_002533755 | South America | Colombia | G35 | G35.C07 | 3103 |
| Hp_002952335 | North America | United States of America | G22 | G22.C04 | 3020 | Hp_003640025 | Europe | Spain | NA | 0.C19 | - |
| Hp_002204015 | Asia | China | G2 | G2.C40 | - | Hp_009740755 | North America | United States of America | G13 | NA | - |
| Hp_006439955 | Asia | Cambodia | G2 | G2.C03 | - | Hp_011174945 | North America | Nicaragua | G37 | G37.C17 | - |
| Hp_002013065 | South America | Colombia | G35 | G35.C07 | 3091 | Hp_002027045 | North America | Mexico | G30 | G30.C01 | 3550 |
| Hp_000591395 | Asia | China | G2 | G2.C43 | 2562 | Hp_006439815 | Asia | Cambodia | G2 | G2.C09 | - |
| Hp_004013055 | Europe | Switzerland | G16 | G16.C01 | - | Hp_002534235 | South America | Colombia | NA | 0.C16 | 3052 |
| Hp_009690355 | Asia | China | G2 | G2.C17 | - | Hp_002532625 | South America | Colombia | G35 | G35.C07 | - |
| Hp_000148915 | South America | Peru | G1 | G1.C02 | 3043 | Hp_003059145 | Europe | United Kingdom | G19 | NA | - |
| Hp_006439685 | Asia | Cambodia | NA | NA | - | Hp_001998405 | South America | Colombia | G37 | G37.C08 | 3499 |
| Hp_006440245 | Asia | Cambodia | G2 | G2.C18 | - | Hp_003058825 | Europe | Sweden | NA | 0.C04 | - |
| Hp_003050665 | Oceania | Australia | G22 | G22.C02 | - | Hp_009740385 | North America | United States of America | G37 | G37.C20 | - |
| Hp_004012315 | Europe | Switzerland | G31 | NA | - | Hp_004013205 | Europe | Switzerland | G15 | G15.C02 | - |
| Hp_003636585 | Oceania | Australia | G24 | G24.C01 | - | Hp_003635795 | Oceania | Australia | G6 | G6.C01 | - |
| Hp_003059545 | Europe | France | NA | 0.C11 | - | Hp_011141055 | North America | Mexico | G25 | NA | - |
| Hp_000317875 | North America | Canada | G1 | G1.C02 | 2945 | Hp_003636205 | Oceania | Australia | G20 | G20.C07 | 1889 |
| Hp_003639305 | Europe | Spain | G26 | G26.C03 | - | Hp_002012405 | South America | Colombia | G37 | G37.C03 | 3512 |
| Hp_003636215 | Oceania | Australia | G20 | G20.C04 | 664 | Hp_002013145 | South America | Colombia | G35 | G35.C07 | 3118 |
| Hp_003059065 | Europe | Sweden | G20 | NA | 1233 | Hp_009741085 | North America | United States of America | G37 | NA | - |
| Hp_002012415 | South America | Colombia | G26 | G26.C04 | 3092 | Hp_002026595 | North America | Mexico | G30 | G30.C01 | 3531 |
| Hp_003635765 | Oceania | Australia | G2 | G2.C10 | - | Hp_002357475 | Asia | Japan | G2 | G2.C13 | - |
| Hp_003059715 | Europe | France | G27 | G27.C01 | - | Hp_003638085 | Oceania | Australia | G29 | NA | - |
| Hp_009690185 | Asia | China | G2 | G2.C17 | - | Hp_002532745 | South America | Colombia | G35 | G35.C07 | 3061 |
| Hp_001998295 | South America | Colombia | G35 | G35.C05 | 3506 | Hp_006438475 | Asia | Cambodia | G2 | G2.C03 | - |
| Hp_001278515 | Asia | Kuwait | G25 | NA | - | Hp_008681225 | North America | Mexico | G37 | G37.C13 | - |
| Hp_002093735 | Africa | Morocco | G37 | G37.C25 | 3496 | Hp_006337365 | Europe | Germany | G25 | G25.C11 | - |
| Hp_000256075 | Asia | India | G3 | NA | - | Hp_003059365 | Europe | France | G25 | G25.C03 | - |
| Hp_004013495 | Europe | Switzerland | G24 | G24.C02 | - | Hp_004012255 | Europe | Switzerland | G7 | G7.C03 | - |
| Hp_000287775 | Asia | China | G2 | G2.C19 | 3674 | Hp_006438525 | Asia | Cambodia | G5 | G5.C01 | - |
| Hp_001189655 | Asia | Malaysia | G25 | G25.C07 | - | Hp_002012755 | South America | Colombia | G34 | G34.C02 | 3140 |
| Hp_009690205 | Asia | China | G2 | G2.C20 | - | Hp_000448405 | South America | Colombia | G37 | G37.C23 | 3510 |
| Hp_000296055 | Asia | Malaysia | G2 | G2.C21 | 3671 | Hp_003059455 | Europe | France | G7 | G7.C02 | - |
| Hp_000470355 | Europe | Germany | G14 | G14.C02 | - | Hp_003636355 | Oceania | Australia | G3 | G3.C04 | - |
| Hp_006439845 | Asia | Cambodia | G4 | G4.C02 | - | Hp_004012765 | Europe | Switzerland | G9 | G9.C01 | - |
| Hp_004323225 | Asia | Malaysia | G2 | G2.C22 | - | Hp_003058785 | Europe | United Kingdom | G9 | NA | - |
| Hp_000444325 | Asia | Malaysia | G2 | G2.C20 | 3679 | Hp_002026485 | North America | Mexico | G30 | G30.C01 | 3530 |
| Hp_003635595 | Oceania | Australia | G20 | G20.C01 | 1258 | Hp_003639795 | Europe | Spain | NA | 0.C09 | - |
| Hp_002951975 | North America | United States of America | G22 | G22.C04 | - | Hp_002027005 | North America | Mexico | G36 | G36.C04 | 3546 |
| Hp_002533415 | South America | Colombia | G35 | G35.C05 | 3123 | Hp_002357615 | Asia | Japan | G2 | G2.C13 | - |
| Hp_002831995 | Asia | India | G11 | G11.C04 | - | Hp_000277425 | South America | Peru | G37 | NA | 3028 |
| Hp_004016925 | Europe | Switzerland | G25 | NA | - | Hp_004014435 | Europe | Switzerland | G29 | G29.C03 | - |
| Hp_001484135 | Asia | Malaysia | G2 | G2.C24 | - | Hp_011175045 | North America | Nicaragua | G24 | G24.C03 | - |
| Hp_003637465 | Oceania | Australia | G7 | NA | - | Hp_004013465 | Europe | Switzerland | G25 | NA | - |
| Hp_002013115 | South America | Colombia | G35 | G35.C07 | 3515 | Hp_001189775 | Asia | Malaysia | G3 | G3.C04 | - |
| Hp_002952475 | North America | United States of America | G22 | G22.C03 | 3020 | Hp_011175195 | North America | Nicaragua | G33 | G33.C01 | - |
| Hp_004117965 | Asia | China(Taiwan) | NA | NA | - | Hp_004323375 | Asia | Malaysia | G2 | G2.C47 | 3684 |
| Hp_004014075 | Europe | Switzerland | G10 | G10.C03 | - | Hp_002357335 | Asia | Japan | G2 | G2.C39 | - |
| Hp_003639485 | Europe | Spain | NA | NA | - | Hp_003638295 | Oceania | Australia | G37 | G37.C30 | 1213 |
| Hp_003059505 | Europe | France | G37 | NA | 2861 | Hp_002832355 | Asia | India | G11 | NA | - |
| Hp_004013975 | Europe | Switzerland | NA | 0.C12 | - | Hp_006337625 | Europe | Germany | G9 | G9.C02 | - |
| Hp_003815575 | South America | Colombia | NA | 0.C16 | 3052 | Hp_006440055 | Asia | Cambodia | G2 | NA | - |
| Hp_002532715 | South America | Colombia | G35 | G35.C07 | 3095 | Hp_009740405 | North America | United States of America | G24 | G24.C03 | - |
| Hp_003060085 | Europe | France | G20 | G20.C01 | - | Hp_000513515 | Oceania | Australia | G20 | G20.C03 | 1404 |
| Hp_004013275 | Europe | Switzerland | G17 | G17.C02 | - | Hp_011140825 | North America | Mexico | NA | 0.C18 | - |
| Hp_000444785 | Asia | Malaysia | G3 | G3.C14 | 3683 | Hp_006440215 | Asia | Cambodia | NA | 0.C01 | - |
| Hp_004014595 | Europe | Switzerland | G37 | G37.C21 | - | Hp_011175315 | North America | Nicaragua | G37 | G37.C10 | - |
| Hp_003639805 | Europe | Spain | G33 | NA | - | Hp_003059105 | Europe | Sweden | G17 | G17.C01 | - |
| Hp_004013135 | Europe | Switzerland | G25 | G25.C09 | - | Hp_001998385 | South America | Colombia | G37 | G37.C02 | 3509 |
| Hp_001248015 | Asia | Malaysia | G3 | G3.C04 | - | Hp_002026925 | North America | Mexico | G35 | G35.C02 | 3526 |
| Hp_006338815 | Europe | Germany | G9 | G9.C02 | - | Hp_001542385 | Asia | Malaysia | G2 | G2.C08 | - |
| Hp_000469985 | Europe | Germany | G14 | G14.C02 | - | Hp_006337745 | Europe | Germany | G18 | G18.C01 | - |
| Hp_004012375 | Europe | Switzerland | G25 | G25.C05 | - | Hp_003636785 | Oceania | Australia | G2 | G2.C27 | - |
| Hp_000448525 | Africa | South Africa | G28 | G28.C02 | 3583 | Hp_006337285 | Europe | Germany | G9 | G9.C02 | - |
| Hp_000689035 | North America | United States of America | G5 | G5.C02 | 204 | Hp_003059285 | Europe | France | G25 | G25.C03 | - |
| Hp_005796135 | Europe | France | G17 | G17.C01 | 3655 | Hp_003059985 | Europe | France | G23 | G23.C01 | - |
| Hp_001185095 | Asia | Malaysia | G3 | G3.C14 | - | Hp_000590775 | Africa | South Africa | G28 | G28.C02 | 3582 |
| Hp_002012475 | South America | Colombia | G35 | G35.C05 | 3107 | Hp_002534395 | South America | Colombia | G37 | G37.C09 | 3087 |
| Hp_003636275 | Oceania | Australia | NA | 0.C11 | - | Hp_011058095 | South America | Colombia | G35 | G35.C06 | - |
| Hp_002533335 | South America | Colombia | G35 | G35.C06 | 3127 | Hp_003636665 | Oceania | Australia | G2 | G2.C19 | - |
| Hp_008033035 | North America | Mexico | G31 | NA | - | Hp_002832325 | Asia | India | G8 | G8.C02 | - |
| Hp_002534455 | South America | Colombia | G35 | G35.C07 | 3086 | Hp_000470135 | Europe | Germany | G18 | G18.C01 | - |
| Hp_000688955 | North America | United States of America | G5 | G5.C02 | - | Hp_011141135 | North America | Mexico | G37 | G37.C06 | - |
| Hp_001653415 | Africa | South Africa | G37 | G37.C18 | 340 | Hp_006440025 | Asia | Cambodia | G5 | NA | - |
| Hp_008681315 | North America | Mexico | G33 | G33.C02 | - | Hp_002191215 | Oceania | Australia | G20 | G20.C03 | 2335 |
| Hp_002532895 | South America | Colombia | G35 | G35.C05 | 3082 | Hp_002357415 | Asia | Japan | G2 | G2.C31 | - |
| Hp_002534515 | South America | Colombia | G37 | G37.C02 | 3067 | Hp_004014195 | Europe | Switzerland | G14 | NA | - |
| Hp_004014305 | Europe | Switzerland | G29 | G29.C03 | - | Hp_004013945 | Europe | Switzerland | G2 | G2.C20 | - |
| Hp_002533615 | South America | Colombia | G26 | G26.C04 | 3112 | Hp_006438415 | Asia | Cambodia | NA | 0.C02 | - |
| Hp_000591235 | Asia | China | G2 | G2.C42 | 2566 | Hp_004014045 | Europe | Switzerland | G24 | G24.C01 | - |
| Hp_002952675 | North America | United States of America | G22 | G22.C04 | 3020 | Hp_004013665 | Europe | Switzerland | NA | NA | - |
| Hp_001940115 | Asia | Singapore | G2 | G2.C15 | - | Hp_002026955 | North America | Mexico | G30 | G30.C01 | 3540 |
| Hp_011175075 | North America | Nicaragua | G37 | G37.C10 | - | Hp_003637695 | Oceania | Australia | G25 | G25.C08 | - |
| Hp_003637665 | Oceania | Australia | G28 | NA | - | Hp_002012725 | South America | Colombia | G35 | G35.C05 | 3097 |
| Hp_003635845 | Oceania | Australia | G15 | G15.C02 | - | Hp_009740605 | North America | United States of America | G33 | G33.C02 | - |
| Hp_000499325 | Europe | Russia | G2 | G2.C01 | - | Hp_006337425 | Europe | Germany | G9 | G9.C02 | - |
| Hp_002832275 | Asia | India | G3 | G3.C11 | - | Hp_003637135 | Oceania | Australia | G4 | G4.C02 | - |
| Hp_011140735 | North America | Mexico | G30 | G30.C01 | 3530 | Hp_003636325 | Oceania | Australia | G37 | G37.C16 | - |
| Hp_000591175 | Asia | China | G2 | G2.C43 | 2562 | Hp_002533605 | South America | Colombia | G35 | G35.C07 | - |
| Hp_011140915 | North America | Mexico | G37 | G37.C11 | 3528 | Hp_003059425 | Europe | France | NA | 0.C11 | - |
| Hp_002533525 | South America | Colombia | G35 | G35.C07 | 3118 | Hp_003638475 | Oceania | Australia | G25 | G25.C01 | - |
| Hp_003639685 | Europe | Spain | G28 | G28.C01 | - | Hp_000444495 | Asia | Malaysia | G3 | NA | 3686 |
| Hp_002534105 | South America | Colombia | G35 | G35.C07 | 3072 | Hp_004012225 | Europe | Switzerland | G17 | G17.C02 | - |
| Hp_002832135 | Asia | India | G8 | G8.C02 | - | Hp_002357555 | Asia | Japan | G2 | G2.C34 | - |
| Hp_002831835 | Asia | India | G3 | G3.C07 | - | Hp_002026805 | North America | Mexico | G37 | G37.C11 | 3545 |

| | | | | | |
|---|---|---|---|---|---|
| Hp_009740525 | North America | United States of America | G37 | NA | - |
| Hp_000224535 | South America | Peru | G1 | G1.C01 | - |
| Hp_004012435 | Europe | Switzerland | G25 | G25.C09 | - |
| Hp_004013025 | Europe | Switzerland | G2 | G2.C35 | - |
| Hp_004323305 | Oceania | Australia | G6 | NA | - |
| Hp_003639845 | Europe | Spain | G12 | G12.C01 | - |
| Hp_009689945 | Asia | China | G2 | G2.C30 | - |
| Hp_901687245 | Oceania | Australia | G9 | G9.C04 | - |
| Hp_000262655 | Asia | China | G2 | NA | 3039 |
| Hp_000966875 | North America | United States of America | G22 | G22.C04 | 3020 |
| Hp_004014255 | Europe | Switzerland | G1 | NA | - |
| Hp_002013175 | South America | Colombia | G35 | G35.C07 | 3122 |
| Hp_002952415 | North America | United States of America | G22 | G22.C03 | 3020 |
| Hp_001446295 | Asia | Malaysia | G2 | G2.C14 | - |
| Hp_003815515 | South America | Colombia | G37 | G37.C08 | 2958 |
| Hp_000401395 | Asia | Malaysia | G11 | G11.C02 | - |
| Hp_003636025 | Oceania | Australia | NA | 0.C03 | - |
| Hp_002357755 | Asia | Japan | G2 | G2.C33 | - |
| Hp_011105755 | North America | Mexico | G31 | G31.C05 | 3534 |
| Hp_002952555 | North America | United States of America | G22 | G22.C03 | 3020 |
| Hp_000296135 | Asia | Malaysia | G2 | G2.C29 | - |
| Hp_002534145 | South America | Colombia | G35 | G35.C07 | 3069 |
| Hp_002832025 | Asia | India | G11 | G11.C04 | - |
| Hp_006440325 | Asia | Cambodia | G4 | G4.C02 | - |
| Hp_011175225 | North America | Nicaragua | G33 | G33.C01 | - |
| Hp_002534095 | South America | Colombia | G35 | G35.C05 | 3073 |
| Hp_011174975 | North America | Nicaragua | G33 | G33.C01 | - |
| Hp_002533995 | South America | Colombia | G35 | G35.C07 | 3079 |
| Hp_002012615 | South America | Colombia | NA | 0.C19 | 3131 |
| Hp_006337875 | Europe | Germany | G23 | G23.C04 | - |
| Hp_002357535 | Asia | Japan | G2 | G2.C32 | - |
| Hp_011140675 | North America | Mexico | G36 | G36.C02 | - |
| Hp_001939585 | Asia | Singapore | G2 | G2.C48 | - |
| Hp_004013415 | Europe | Switzerland | G25 | G25.C10 | - |
| Hp_006438535 | Asia | Cambodia | G2 | G2.C08 | - |
| Hp_003636415 | Oceania | Australia | G24 | G24.C07 | 1380 |
| Hp_009737135 | North America | United States of America | G20 | G20.C03 | 2335 |
| Hp_009740975 | North America | United States of America | G37 | G37.C28 | - |
| Hp_006338545 | Europe | Germany | G18 | G18.C01 | - |
| Hp_002533975 | South America | Colombia | G35 | NA | - |
| Hp_003059195 | Europe | Belgium | G9 | G9.C04 | - |
| Hp_003059565 | Europe | France | G37 | G37.C24 | - |
| Hp_003636555 | Oceania | Australia | G2 | G2.C03 | - |
| Hp_009689755 | Asia | China | G2 | G2.C35 | - |
| Hp_002533495 | South America | Colombia | G35 | G35.C05 | 3117 |
| Hp_003636085 | Oceania | Australia | G2 | G2.C10 | - |
| Hp_001549875 | Asia | China(Taiwan) | G2 | G2.C38 | - |
| Hp_003637405 | Oceania | Australia | G25 | G25.C10 | - |
| Hp_003059045 | Europe | Sweden | G20 | NA | - |
| Hp_009741205 | North America | United States of America | G37 | G37.C22 | - |
| Hp_009740495 | North America | United States of America | G37 | G37.C29 | - |
| Hp_002831915 | Asia | India | G3 | G3.C10 | - |
| Hp_004013555 | Europe | Switzerland | G24 | NA | - |
| Hp_006439655 | Asia | Cambodia | G2 | G2.C03 | - |
| Hp_002026545 | North America | Mexico | G37 | G37.C11 | 3529 |
| Hp_002026935 | North America | Mexico | G37 | G37.C11 | 3528 |
| Hp_006440265 | Asia | Cambodia | G2 | G2.C40 | - |
| Hp_002532795 | South America | Colombia | NA | 0.C16 | 3062 |
| Hp_000401375 | Asia | Malaysia | G2 | G2.C18 | - |
| Hp_002013045 | South America | Colombia | G26 | G26.C04 | 3514 |
| Hp_002832415 | Asia | India | G11 | NA | - |
| Hp_004012115 | Europe | Switzerland | G17 | G17.C02 | - |
| Hp_000766035 | Asia | Indonesia | G2 | G2.C15 | - |
| Hp_002533055 | South America | Colombia | G35 | G35.C05 | 3139 |
| Hp_003639285 | Europe | Spain | G32 | G32.C02 | - |
| Hp_003637755 | Oceania | Australia | G37 | G37.C33 | - |
| Hp_003638355 | Oceania | Australia | G24 | G24.C03 | - |
| Hp_011140555 | North America | Mexico | G36 | G36.C04 | - |
| Hp_003639745 | Europe | Spain | NA | 0.C16 | - |
| Hp_001189685 | Asia | Malaysia | G2 | G2.C18 | - |
| Hp_002832295 | Asia | India | G8 | G8.C01 | - |
| Hp_000591195 | Asia | China | G2 | G2.C42 | 2566 |
| Hp_011175145 | North America | Nicaragua | G37 | G37.C17 | - |
| Hp_003638465 | Oceania | Australia | G23 | G23.C01 | - |
| Hp_003637265 | Oceania | Australia | NA | NA | - |
| Hp_002534225 | South America | Colombia | G35 | G35.C07 | 3064 |
| Hp_002952275 | North America | United States of America | G22 | G22.C04 | 3020 |
| Hp_003635705 | Oceania | Australia | G29 | G29.C04 | - |
| Hp_004013295 | Europe | Switzerland | G25 | NA | - |
| Hp_003639455 | Europe | Spain | G25 | G25.C06 | - |
| Hp_003638245 | Oceania | Australia | G20 | G20.C04 | 1145 |
| Hp_004014575 | Europe | Switzerland | NA | NA | - |
| Hp_006440155 | Asia | Cambodia | G2 | G2.C04 | - |
| Hp_011140415 | North America | Mexico | G31 | G31.C01 | 3523 |
| Hp_003639365 | Europe | Spain | G32 | G32.C01 | - |
| Hp_004117945 | Asia | China(Taiwan) | G2 | G2.C45 | - |
| Hp_000591315 | Asia | China | G2 | G2.C43 | 2562 |
| Hp_001542495 | Asia | Malaysia | G2 | G2.C40 | - |
| Hp_000444725 | Asia | Malaysia | G2 | G2.C47 | 3684 |
| Hp_003636675 | Oceania | Australia | G2 | G2.C08 | - |
| Hp_002026505 | North America | Mexico | G37 | G37.C11 | 3522 |
| Hp_009690405 | Asia | China | G2 | G2.C21 | - |
| Hp_002357855 | Asia | Japan | G2 | G2.C13 | - |
| Hp_002012495 | South America | Colombia | G26 | G26.C03 | 3094 |
| Hp_008681235 | North America | Mexico | G30 | G30.C01 | - |
| Hp_006440125 | Asia | Cambodia | G2 | G2.C03 | - |
| Hp_011141035 | North America | Mexico | G30 | G30.C01 | - |
| Hp_003636995 | Oceania | Australia | NA | 0.C11 | - |
| Hp_003639425 | Europe | Spain | NA | 0.C09 | - |
| Hp_003060025 | Europe | France | G29 | G29.C01 | - |
| Hp_001248105 | Asia | Malaysia | G3 | NA | - |
| Hp_002831975 | Asia | India | G11 | NA | - |
| Hp_004014145 | Europe | Switzerland | G24 | G24.C06 | - |
| Hp_000277365 | South America | Peru | G1 | G1.C02 | 3046 |
| Hp_002012645 | South America | Colombia | G26 | G26.C04 | 3112 |
| Hp_008326525 | Asia | Japan | G2 | G2.C23 | - |
| Hp_004012955 | Europe | Switzerland | G25 | NA | - |
| Hp_006338685 | Europe | Germany | G20 | G20.C02 | - |
| Hp_004013325 | Europe | Switzerland | G14 | NA | - |
| Hp_002012965 | South America | Colombia | G35 | G35.C07 | 3093 |
| Hp_000730525 | South America | Colombia | G35 | G35.C05 | 3097 |
| Hp_006439875 | Asia | Cambodia | G4 | G4.C02 | - |
| Hp_002532985 | South America | Colombia | G35 | G35.C07 | 3104 |
| Hp_004323315 | Oceania | Australia | G21 | G21.C01 | - |
| Hp_001484185 | Asia | Malaysia | G3 | G3.C14 | - |
| Hp_006439795 | Asia | Cambodia | G4 | G4.C02 | - |
| Hp_011058105 | South America | Colombia | G26 | G26.C04 | - |
| Hp_011175375 | North America | Nicaragua | G33 | G33.C01 | - |
| Hp_003636905 | Oceania | Australia | G2 | G2.C17 | - |
| Hp_003060035 | Europe | France | G15 | G15.C01 | - |
| Hp_002012885 | South America | Colombia | G35 | G35.C05 | - |
| Hp_009798365 | Africa | Morocco | G37 | G37.C25 | 3496 |
| Hp_004014005 | Europe | Switzerland | NA | 0.C07 | - |
| Hp_002357835 | Asia | Japan | G2 | G2.C33 | - |
| Hp_004013905 | Europe | Switzerland | G9 | G9.C01 | - |
| Hp_003638525 | Oceania | Australia | G19 | G19.C03 | 975 |
| Hp_009689695 | Asia | China | G2 | G2.C44 | - |
| Hp_001051485 | Asia | Malaysia | G3 | NA | - |
| Hp_002534545 | South America | Colombia | G35 | G35.C07 | 3055 |
| Hp_008681485 | North America | Mexico | G37 | G37.C12 | - |
| Hp_001999845 | South America | Colombia | G35 | G35.C06 | 3497 |
| Hp_003637025 | Oceania | Australia | G2 | G2.C20 | - |
| Hp_009740965 | North America | United States of America | NA | 0.C14 | - |
| Hp_001998125 | South America | Colombia | G37 | G37.C04 | 3498 |
| Hp_001976105 | Asia | Malaysia | G3 | G3.C14 | - |
| Hp_002534065 | South America | Colombia | NA | 0.C16 | 3075 |
| Hp_009740645 | North America | United States of America | G37 | G37.C23 | - |
| Hp_003059865 | Europe | France | G29 | G29.C01 | - |
| Hp_009690135 | Asia | China | G2 | G2.C08 | - |
| Hp_003640585 | Asia | Vietnam | G2 | NA | 3170 |
| Hp_002357675 | Asia | Japan | G2 | G2.C34 | - |
| Hp_001446385 | Asia | Malaysia | G2 | G2.C18 | - |
| Hp_009740725 | North America | United States of America | G37 | G37.C26 | - |
| Hp_001940145 | Asia | Singapore | G2 | G2.C48 | - |
| Hp_003640055 | Europe | Spain | G27 | G27.C01 | - |
| Hp_003058905 | Europe | France | G12 | G12.C02 | - |
| Hp_011140665 | North America | Mexico | G31 | G31.C05 | - |
| Hp_009690275 | Asia | China | G2 | NA | - |
| Hp_003636855 | Oceania | Australia | G29 | NA | - |
| Hp_011140535 | North America | Mexico | G31 | G31.C05 | - |
| Hp_001940095 | Asia | Singapore | G2 | G2.C46 | - |
| Hp_004013405 | Europe | Switzerland | G17 | G17.C02 | - |
| Hp_004012145 | Europe | Switzerland | G17 | G17.C02 | - |
| Hp_001446325 | Asia | Malaysia | G2 | G2.C45 | - |
| Hp_009740785 | North America | United States of America | G37 | G37.C15 | - |
| Hp_001998205 | South America | Colombia | G37 | NA | 3502 |
| Hp_000688935 | North America | United States of America | G5 | G5.C02 | - |
| Hp_002027065 | North America | Mexico | G37 | G37.C11 | 3544 |
| Hp_006338145 | Europe | Germany | G24 | G24.C04 | - |
| Hp_002357695 | Asia | Japan | G2 | G2.C16 | - |
| Hp_000448465 | South America | Colombia | G37 | G37.C04 | - |
| Hp_004012705 | Europe | Switzerland | G18 | G18.C02 | - |
| Hp_008681425 | North America | Mexico | G35 | G35.C06 | - |
| Hp_002900175 | Europe | Germany | G9 | G9.C04 | - |
| Hp_002831855 | Asia | India | G3 | G3.C06 | - |
| Hp_002832155 | Asia | India | G7 | NA | - |
| Hp_003059305 | Europe | France | G37 | G37.C32 | - |
| Hp_000591255 | Asia | China | G2 | G2.C43 | 2563 |
| Hp_006337305 | Europe | Germany | G24 | G24.C08 | - |
| Hp_011140755 | North America | Mexico | NA | 0.C17 | - |
| Hp_009689835 | Asia | China | G2 | G2.C37 | - |
| Hp_003639935 | Europe | Spain | G32 | G32.C02 | - |
| Hp_003639545 | Europe | Spain | G35 | NA | - |
| Hp_004323235 | Asia | Malaysia | G6 | G6.C01 | - |
| Hp_011175115 | North America | Nicaragua | G33 | G33.C01 | - |
| Hp_006439775 | Asia | Cambodia | G4 | G4.C02 | - |
| Hp_004013155 | Europe | Switzerland | G9 | G9.C05 | - |
| Hp_003637415 | Oceania | Australia | G29 | G29.C02 | - |
| Hp_011175395 | North America | Nicaragua | G37 | NA | - |
| Hp_003638585 | Oceania | Australia | G20 | G20.C04 | 1143 |
| Hp_003638055 | Oceania | Australia | G6 | NA | - |
| Hp_002026705 | North America | Mexico | G37 | G37.C07 | 3537 |
| Hp_000020245 | South America | Peru | G1 | G1.C02 | 3047 |
| Hp_002532925 | South America | Colombia | G35 | G35.C05 | 3084 |
| Hp_008033055 | North America | Mexico | G37 | G37.C07 | 3537 |
| Hp_002533355 | South America | Colombia | G37 | G37.C08 | 3125 |
| Hp_002534435 | South America | Colombia | G35 | G35.C07 | 3089 |
| Hp_003059165 | Europe | Belgium | NA | NA | - |
| Hp_002832405 | Asia | India | G7 | G7.C02 | - |
| Hp_000689055 | North America | United States of America | G5 | G5.C02 | 204 |
| Hp_002533465 | South America | Colombia | G35 | G35.C07 | 3121 |
| Hp_003636155 | Oceania | Australia | G15 | G15.C02 | - |
| Hp_000392535 | Asia | Malaysia | G2 | G2.C25 | 3682 |
| Hp_009740465 | North America | United States of America | G37 | G37.C20 | - |
| Hp_006440415 | Asia | Cambodia | G5 | G5.C01 | - |
| Hp_001998245 | South America | Colombia | G35 | G35.C05 | 3503 |
| Hp_002952535 | North America | United States of America | G22 | G22.C04 | 3020 |
| Hp_002026455 | North America | Mexico | NA | 0.C14 | 3527 |
| Hp_002533205 | South America | Colombia | G26 | G26.C04 | 3132 |
| Hp_003637295 | Oceania | Australia | G25 | NA | - |
| Hp_002357735 | Asia | Japan | G2 | G2.C34 | - |
| Hp_002005525 | Oceania | Australia | G9 | G9.C04 | - |
| Hp_006338285 | Europe | Germany | G21 | G21.C01 | - |
| Hp_006337345 | Europe | Germany | G9 | G9.C02 | - |
| Hp_003059485 | Europe | France | G12 | NA | - |
| Hp_004012515 | Europe | Switzerland | G16 | G16.C01 | - |

| | | | | | |
|---|---|---|---|---|---|
| Hp_001997995 | South America | Colombia | G37 | NA | 3505 |
| Hp_002952495 | North America | United States of America | G22 | G22.C03 | 3020 |
| Hp_009689605 | Asia | China | G2 | G2.C05 | - |
| Hp_003638535 | Oceania | Australia | G9 | G9.C05 | - |
| Hp_011140335 | North America | Mexico | G28 | G28.C01 | - |
| Hp_004014095 | Europe | Switzerland | G23 | G23.C03 | - |
| Hp_000401315 | Asia | Malaysia | G3 | NA | - |
| Hp_004323455 | Oceania | Australia | G37 | G37.C16 | - |
| Hp_003815595 | South America | Colombia | G35 | G35.C07 | 3105 |
| Hp_002832475 | Asia | India | G3 | G3.C13 | - |
| Hp_003637965 | Oceania | Australia | G21 | G21.C02 | - |
| Hp_008032955 | North America | Mexico | NA | 0.C18 | - |
| Hp_002533025 | South America | Colombia | G35 | G35.C07 | 3105 |
| Hp_004323515 | Oceania | Australia | G10 | G10.C01 | - |
| Hp_001189765 | Asia | Malaysia | G2 | G2.C22 | - |
| Hp_011140615 | North America | Mexico | G36 | G36.C01 | - |
| Hp_000444385 | Asia | Malaysia | G3 | G3.C10 | - |
| Hp_003639985 | Europe | Spain | NA | 0.C06 | - |
| Hp_009689885 | Asia | China | G2 | G2.C30 | - |
| Hp_003058995 | Europe | Sweden | NA | 0.C05 | - |
| Hp_009690125 | Asia | China | G2 | G2.C09 | - |
| Hp_009740735 | North America | United States of America | G37 | G37.C26 | - |
| Hp_004012775 | Europe | Switzerland | G18 | G18.C02 | - |
| Hp_002533915 | South America | Colombia | G35 | G35.C07 | 3091 |
| Hp_004012895 | Europe | Switzerland | G9 | G9.C03 | - |
| Hp_003636525 | Oceania | Australia | NA | 0.C03 | - |
| Hp_006439745 | Asia | Cambodia | G2 | G2.C02 | - |
| Hp_002534305 | South America | Colombia | G35 | G35.C05 | 3102 |
| Hp_002357355 | Asia | Japan | G2 | G2.C39 | - |
| Hp_006438465 | Asia | Cambodia | G2 | G2.C03 | - |
| Hp_011140475 | North America | Mexico | NA | 0.C08 | - |
| Hp_004014285 | Europe | Switzerland | G25 | NA | - |
| Hp_004088075 | Asia | China | G2 | G2.C19 | - |
| Hp_000591375 | Asia | China | G2 | G2.C43 | 2561 |
| Hp_004014625 | Europe | Switzerland | G10 | G10.C03 | - |
| Hp_003637885 | Oceania | Australia | G37 | G37.C33 | - |
| Hp_003636615 | Oceania | Australia | G16 | G16.C03 | 3255 |
| Hp_001185015 | Asia | Malaysia | G6 | G6.C01 | - |
| Hp_004013525 | Europe | Switzerland | G25 | G25.C05 | - |
| Hp_009689965 | Asia | China | G2 | G2.C07 | - |
| Hp_003639865 | Europe | Spain | NA | 0.C08 | - |
| Hp_004014105 | Europe | Switzerland | G7 | G7.C01 | - |
| Hp_003635655 | Oceania | Australia | G13 | G13.C02 | - |
| Hp_008681395 | North America | Mexico | G35 | G35.C06 | - |
| Hp_000451025 | Asia | Malaysia | G2 | G2.C25 | 3682 |
| Hp_002532815 | South America | Colombia | G35 | G35.C05 | - |
| Hp_002533275 | South America | Colombia | NA | 0.C19 | 3131 |
| Hp_002026895 | North America | Mexico | G37 | G37.C01 | 3524 |
| Hp_000333835 | Europe | Russia | NA | NA | 1767 |
| Hp_003636335 | Oceania | Australia | G15 | G15.C02 | - |
| Hp_002533135 | South America | Colombia | G34 | G34.C02 | 3140 |
| Hp_003637605 | Oceania | Australia | G24 | NA | - |
| Hp_002832525 | Asia | India | G25 | G25.C02 | - |
| Hp_002533695 | South America | Colombia | G35 | G35.C05 | 3107 |
| Hp_002012915 | South America | Colombia | G35 | G35.C05 | 3123 |
| Hp_000685665 | Oceania | Australia | G13 | G13.C01 | - |
| Hp_009690965 | Asia | Japan | G2 | G2.C31 | - |
| Hp_003640155 | Europe | Spain | NA | 0.C13 | - |
| Hp_011140995 | North America | Mexico | G34 | G34.C03 | 3520 |
| Hp_002832615 | Asia | India | G8 | G8.C02 | - |
| Hp_003639575 | Europe | Spain | NA | NA | - |
| Hp_003815485 | South America | Colombia | G35 | G35.C05 | - |
| Hp_000295975 | Asia | Malaysia | G3 | G3.C04 | 3669 |
| Hp_002533095 | South America | Colombia | G35 | G35.C07 | 3142 |
| Hp_002012745 | South America | Colombia | G35 | G35.C05 | 3139 |
| Hp_003059385 | Europe | France | G13 | NA | - |
| Hp_006337445 | Europe | Germany | G24 | G24.C08 | - |
| Hp_006337385 | Europe | Germany | G24 | G24.C04 | - |
| Hp_003059445 | Europe | France | G15 | G15.C02 | - |
| Hp_001976125 | Asia | Malaysia | G3 | G3.C14 | - |
| Hp_003636485 | Oceania | Australia | G20 | G20.C01 | - |
| Hp_004012785 | Europe | Switzerland | G9 | G9.C03 | - |
| Hp_000439295 | Asia | Singapore | G2 | G2.C26 | - |
| Hp_002533145 | South America | Colombia | G35 | G35.C05 | 3138 |
| Hp_006339365 | Europe | Germany | G14 | NA | - |
| Hp_004012245 | Europe | Switzerland | G25 | NA | - |
| Hp_000688975 | North America | United States of America | G5 | G5.C02 | - |
| Hp_011175215 | North America | Nicaragua | G37 | NA | - |
| Hp_006440315 | Asia | Cambodia | G2 | G2.C03 | - |
| Hp_001998285 | South America | Colombia | G37 | G37.C08 | 3501 |
| Hp_000591155 | Asia | China | G2 | G2.C43 | 2561 |
| Hp_009740705 | North America | United States of America | G37 | G37.C30 | - |
| Hp_009689875 | Asia | China | G2 | G2.C35 | - |
| Hp_002832255 | Asia | India | G15 | G15.C02 | - |
| Hp_001940035 | Asia | Singapore | G3 | G3.C09 | - |
| Hp_011140595 | North America | Mexico | NA | 0.C15 | 3549 |
| Hp_004323275 | Asia | Malaysia | G3 | G3.C04 | - |
| Hp_009690045 | Asia | China | G2 | G2.C21 | - |
| Hp_002952655 | North America | United States of America | G22 | G22.C04 | 3020 |
| Hp_002533705 | South America | Colombia | G26 | G26.C03 | - |
| Hp_000255955 | North America | El Salvador | NA | NA | 3048 |
| Hp_004013015 | Europe | Switzerland | NA | 0.C11 | - |
| Hp_009690315 | Asia | China | G2 | G2.C36 | - |

| | | | | | |
|---|---|---|---|---|---|
| Hp_003636445 | Oceania | Australia | G25 | G25.C04 | 3153 |
| Hp_003636385 | Oceania | Australia | G29 | G29.C05 | - |
| Hp_006337485 | Europe | Germany | G15 | G15.C01 | - |
| Hp_003058755 | Europe | Sweden | NA | 0.C05 | 958 |
| Hp_005250645 | Europe | Belarus | NA | 0.C05 | - |
| Hp_006439895 | Asia | Cambodia | G4 | G4.C01 | - |
| Hp_001542465 | Asia | Malaysia | G2 | G2.C08 | - |
| Hp_002534375 | South America | Colombia | G26 | G26.C04 | 3514 |
| Hp_004012365 | Europe | Switzerland | G25 | G25.C05 | - |
| Hp_001420425 | Asia | Malaysia | G2 | G2.C45 | - |
| Hp_003637835 | Oceania | Australia | G29 | G29.C02 | - |
| Hp_002026625 | North America | Mexico | G26 | G26.C01 | 3532 |
| Hp_003635545 | Oceania | Australia | G16 | G16.C04 | 1883 |
| Hp_001189815 | Asia | Malaysia | G2 | G2.C44 | - |
| Hp_003635715 | Oceania | Australia | G37 | G37.C33 | - |
| Hp_000224555 | South America | Peru | G1 | G1.C02 | 3042 |
| Hp_008026615 | North America | Mexico | G37 | G37.C11 | 3522 |
| Hp_003636265 | Oceania | Australia | G16 | G16.C03 | - |
| Hp_002533825 | South America | Colombia | G35 | G35.C07 | 3096 |
| Hp_002026515 | North America | Mexico | G37 | G37.C01 | 3521 |
| Hp_003059925 | Europe | France | G12 | G12.C02 | - |
| Hp_900638475 | Africa | Nigeria | G29 | G29.C01 | 1288 |
| Hp_002532595 | South America | Colombia | NA | NA | - |
| Hp_004014175 | Europe | Switzerland | G12 | NA | - |
| Hp_002013015 | South America | Colombia | G35 | G35.C07 | 3096 |
| Hp_004013755 | Europe | Switzerland | NA | NA | - |
| Hp_000258845 | Oceania | Australia | G11 | NA | 415 |
| Hp_002026745 | North America | Mexico | G36 | G36.C01 | 3539 |
| Hp_006439715 | Asia | Cambodia | G2 | G2.C02 | - |
| Hp_004014545 | Europe | Switzerland | G24 | G24.C02 | - |
| Hp_009740855 | North America | United States of America | G31 | G31.C03 | - |
| Hp_001997965 | South America | Colombia | G37 | G37.C23 | 3504 |
| Hp_002026985 | North America | Mexico | G37 | G37.C13 | 3542 |
| Hp_002533855 | South America | Colombia | G26 | G26.C03 | 3094 |
| Hp_003636725 | Oceania | Australia | G15 | G15.C01 | - |
| Hp_001433515 | North America | Mexico | NA | 0.C07 | - |
| Hp_003638135 | Oceania | Australia | G29 | G29.C04 | - |
| Hp_002952355 | North America | United States of America | G22 | G22.C04 | 3020 |
| Hp_003059225 | Europe | France | G37 | NA | - |
| Hp_004013965 | Europe | Switzerland | G10 | G10.C01 | - |
| Hp_001185045 | Asia | Malaysia | G3 | NA | - |
| Hp_003636645 | Oceania | Australia | G2 | G2.C11 | - |
| Hp_003636965 | Oceania | Australia | G37 | G37.C18 | - |
| Hp_000689135 | North America | United States of America | G5 | G5.C02 | - |
| Hp_003636035 | Oceania | Australia | G6 | NA | - |
| Hp_009689615 | Asia | China | G2 | G2.C27 | - |
| Hp_004012985 | Europe | Switzerland | G9 | G9.C05 | - |
| Hp_003058965 | Europe | Sweden | NA | 0.C04 | - |
| Hp_002012945 | South America | Colombia | G35 | G35.C07 | 3055 |
| Hp_004012655 | Europe | Switzerland | G24 | G24.C05 | - |
| Hp_002983705 | North America | United States of America | G37 | G37.C25 | 3496 |
| Hp_003637995 | Oceania | Australia | G2 | G2.C08 | - |
| Hp_009740905 | North America | United States of America | G31 | G31.C04 | - |
| Hp_002534005 | South America | Colombia | G37 | G37.C05 | 3077 |
| Hp_011175235 | North America | Nicaragua | G34 | NA | - |
| Hp_006440335 | Asia | Cambodia | G2 | G2.C03 | - |
| Hp_002533905 | South America | Colombia | G26 | G26.C04 | 3092 |
| Hp_011175175 | North America | Nicaragua | G37 | G37.C11 | - |
| Hp_002832645 | Asia | India | G11 | NA | - |
| Hp_001940015 | Asia | Singapore | G2 | G2.C15 | - |
| Hp_009741025 | North America | United States of America | G2 | G2.C23 | - |
| Hp_002534315 | South America | Colombia | G26 | G26.C04 | 3513 |
| Hp_003638025 | Oceania | Australia | G4 | G4.C02 | - |
| Hp_004013605 | Europe | Switzerland | NA | 0.C06 | - |
| Hp_008681515 | North America | Mexico | NA | 0.C17 | - |
| Hp_002222575 | Asia | China | G2 | G2.C05 | - |
| Hp_008681285 | North America | Mexico | G36 | G36.C04 | - |
| Hp_009689995 | Asia | China | G2 | G2.C30 | - |
| Hp_004014505 | Europe | Switzerland | G25 | NA | - |
| Hp_004012865 | Europe | Switzerland | G10 | G10.C01 | - |
| Hp_003058885 | Europe | Belgium | G27 | G27.C02 | - |
| Hp_002532685 | South America | Colombia | G35 | G35.C07 | 3057 |
| Hp_005796125 | Europe | Poland | G17 | G17.C01 | 3655 |
| Hp_002012855 | South America | Colombia | G37 | G37.C08 | 3098 |
| Hp_002357495 | Asia | Japan | G2 | G2.C16 | - |
| Hp_004014115 | Europe | Switzerland | NA | 0.C09 | - |
| Hp_001484155 | Asia | Malaysia | G2 | G2.C24 | - |
| Hp_009740325 | North America | United States of America | G37 | G37.C14 | - |
| Hp_003636605 | Oceania | Australia | G2 | G2.C03 | - |
| Hp_004014395 | Europe | Switzerland | G15 | G15.C02 | - |
| Hp_004323545 | Oceania | Australia | G3 | NA | - |
| Hp_003636885 | Oceania | Australia | G21 | G21.C02 | - |
| Hp_003059845 | Europe | Ireland | G20 | G20.C07 | - |
| Hp_002532805 | South America | Colombia | G35 | G35.C05 | 3070 |
| Hp_003640085 | Europe | Spain | G31 | G31.C02 | - |
| Hp_008681365 | North America | Mexico | G33 | G33.C02 | - |
| Hp_002832605 | Asia | India | G25 | G25.C02 | - |
| Hp_004012575 | Europe | Switzerland | G32 | G32.C01 | - |
| Hp_002026865 | North America | Mexico | NA | 0.C15 | 3549 |
| Hp_002533535 | South America | Colombia | NA | 0.C20 | 3115 |
| Hp_001999865 | South America | Colombia | G37 | G37.C19 | 3507 |
| Hp_009740535 | North America | United States of America | G37 | G37.C26 | - |
| Hp_000296035 | Asia | Malaysia | G2 | G2.C41 | - |
| Hp_008681455 | North America | Mexico | G37 | G37.C12 | - |

Supplementary Table 2. Genomes information for testing

| GenBank assembly accession | Genome submit date | Collected continent | Collected country | Sequencing platform |
|---|---|---|---|---|
| GCF_002206465.1 | 19/01/2018 | North America | United States of America | sequenced by Pacbio & Illumina |
| GCF_002900195.1 | 25/01/2018 | Europe | Germany | sequenced by Illumina |
| GCF_004307235.1 | 31/01/2019 | Aisa | Japan | sequenced by Illumina |
| GCF_004307275.1 | 31/01/2019 | Aisa | Japan | sequenced by Illumina |
| GCF_004307295.1 | 31/01/2019 | Aisa | Japan | sequenced by Illumina |
| GCF_900478295.1 | 18/06/2018 | Oceania | Australia | sequenced by Illumina |
| GCF_016757535.1 | 27/01/2021 | South America | Peru | sequenced by Illumina |
| GCF_016749335.1 | 27/01/2021 | South America | Peru | sequenced by Illumina |
| GCF_016757535.1 | 27/01/2021 | South America | Peru | sequenced by Illumina |
| GCF_013122115.1 | 19/05/2020 | Europe | Russia | sequenced by Illumina |

### User instruction

*Helicobactor pylori* typing tool (HPTT) is a genomic typing method based on whole genome sequencing (WGS) data, which can facilitate genetic population analysis and epidemiological origin tracing. In our *H. pylori* typing website (https://db.cngb.org/HPTT/), two modules are available, including both genetic typing and geographic visualizing of the global *H. pylori* isolates. Here is the website instruction for users.

#### Homepage

You can see the brief introduction of our HPTT on the homepage of our website. Two modules were displayed at this page: the genomic typing tool of *H. pylori* ("QUERY TOOL") and the phylogeographic visualization tool of *H. pylori* ("VISUALIZATION TOOL").

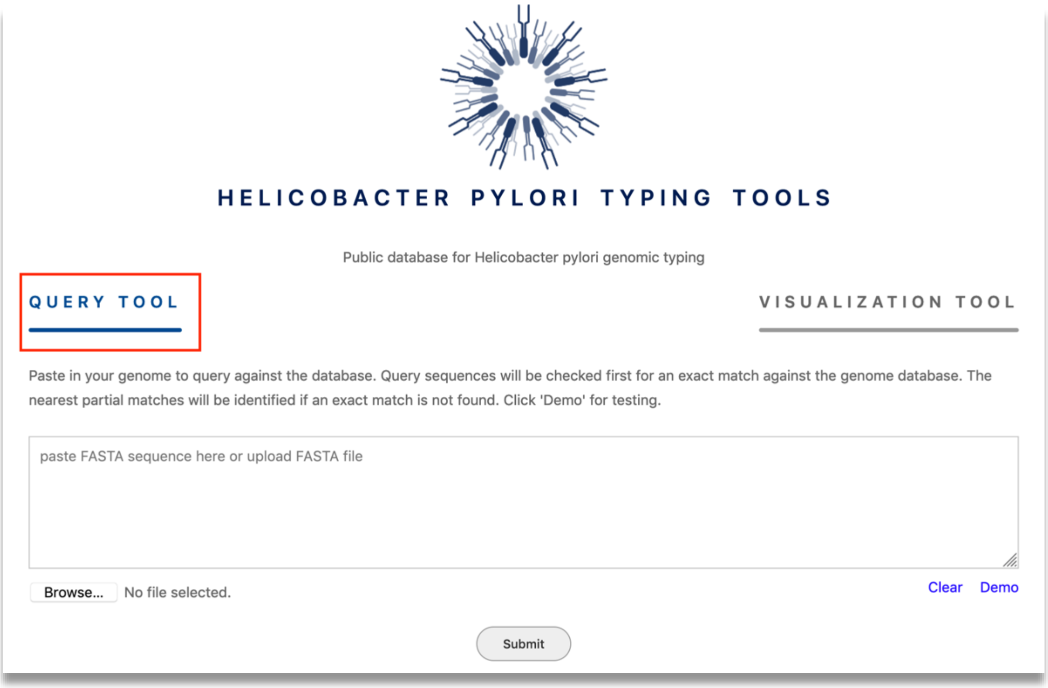

#### - Data compatibility

In order to achieve the best typing results, it is highly recommended to upload sequencing data generated by the MPS (Massive Parallel Sequencing) method. The common MPS platform includes MGI DNBSEQ® platform and Illumina® platform. General workflow for the MPS including sample pre-treatment, library preparation, sequencing and data analysis. Companies who doing MPS would offer the solutions for all steps, including reagent, instrument, etc.

**- The genomic typing tool of *H. pylori***

At the homepage of website, a tested demo genome is provided. Please see the screenshot below. Click the "Demo" button at the bottom right corner of the input box. It may take few seconds since the genome file is relatively large, and there might be a slight freeze during the loading. If the genome is entered incorrectly, click the "Clear" button just next to the "Demo" to clear the input box. For your own sequenced genome, you can either copy and paste in the input box or upload it by click the "Browse" button. After entering/uploading the genome, click the "Submit" button to query against the database.

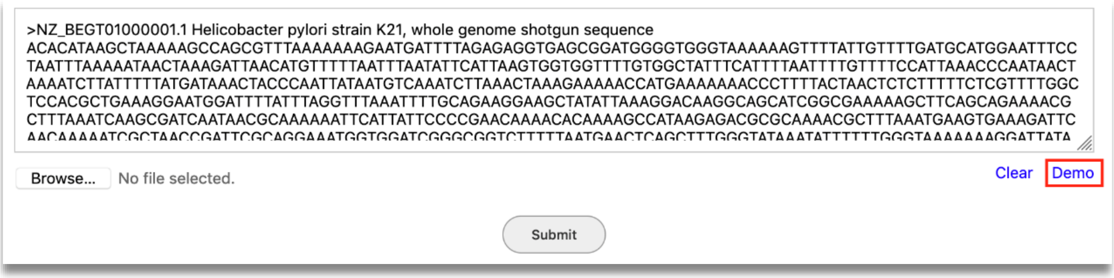

If your uploaded file is neither *H. pylori* nor sequenced by the second-generation sequencing technology, it might be reported as an error. You will need to check whether your genome fit our criteria.

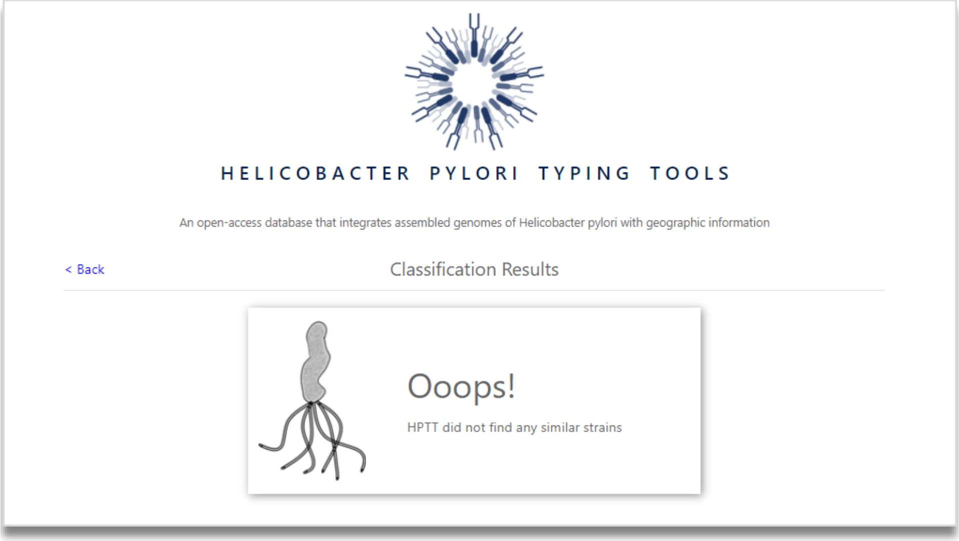

You will get the typing results if it is successfully typed. Please see the image below. Firstly, the strain with the highest identity ("the closest matching strain") that is most similar to our database will be displayed, including region, country, MLST type, data release time, sequenced research institution, etc. Secondly, if the most similar bacterial species has a typing patterns in our database, it will be displayed on the right. If the corresponding types not completely refer to one continent/country of patterns in our database, the proportion of each continent/country will be displayed in different colours in the bar.

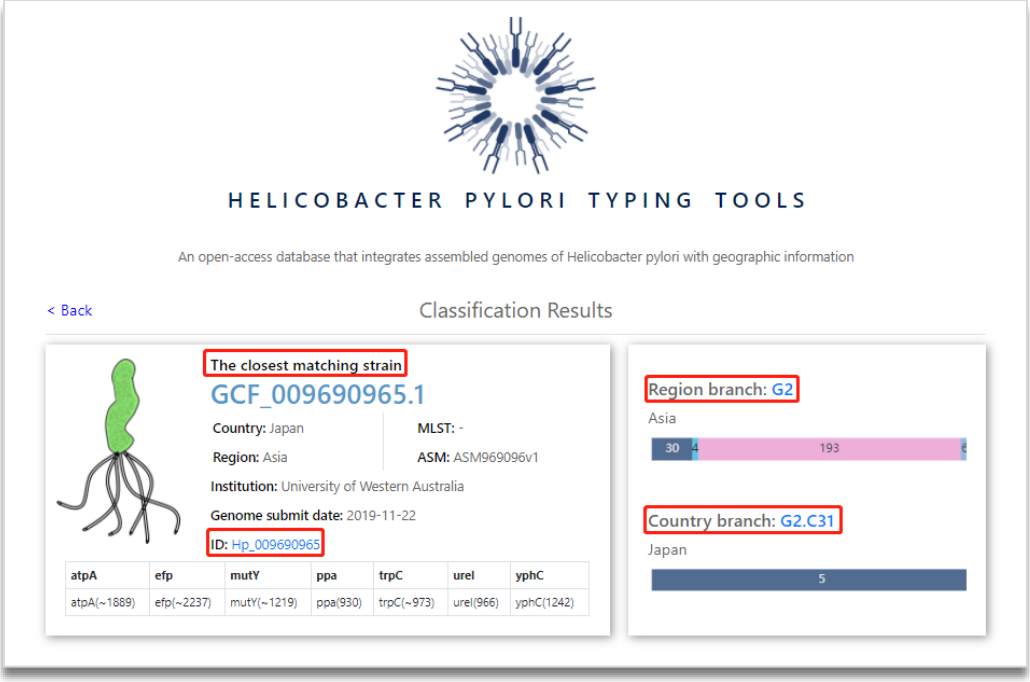

For the region and country groups in our database, each strain can be listed to the corresponding groups. In addition, you can easily access to the visualization tool and NCBI information page by clicking the link (letters in blue).

G2 Region Group List - 233 strains

| Group | Genome_id | Tip_id | Region | Country | Submit_date | MLST | Link(NCBI) |
|---|---|---|---|---|---|---|---|
| G2 | GCF_000262655.1 | Hp_000262655 | Asia | China | 2012-05-14 | 3039 | more |
| G2 | GCF_000287735.1 | Hp_000287735 | Asia | China | 2013-02-23 | 2164 | more |
| G2 | GCF_000287755.1 | Hp_000287755 | Asia | China | 2013-02-23 | 2128 | more |

G2.C31 Country Group List - 5 strains

| Group | Genome_id | Tip_id | Region | Country | Submit_date | MLST | Link(NCBI) |
|---|---|---|---|---|---|---|---|
| G2.C31 | GCF_002357315.1 | Hp_002357315 | Asia | Japan | 2017-01-17 | - | more |
| G2.C31 | GCF_002357415.1 | Hp_002357415 | Asia | Japan | 2017-01-17 | - | more |
| G2.C31 | GCF_002357715.1 | Hp_002357715 | Asia | Japan | 2017-01-17 | - | more |
| G2.C31 | GCF_002357775.1 | Hp_002357775 | Asia | Japan | 2017-01-17 | - | more |
| G2.C31 | GCF_009690965.1 | Hp_009690965 | Asia | Japan | 2019-11-22 | - | more |

**- The visualization tool of *H. pylori***

You can easily access to the visualization tool at the homepage by clicking the "VISUALIZATION TOOL" button. All the genomes in our database can be interacted and displayed in both phylogenetic relationships and geographic locations.

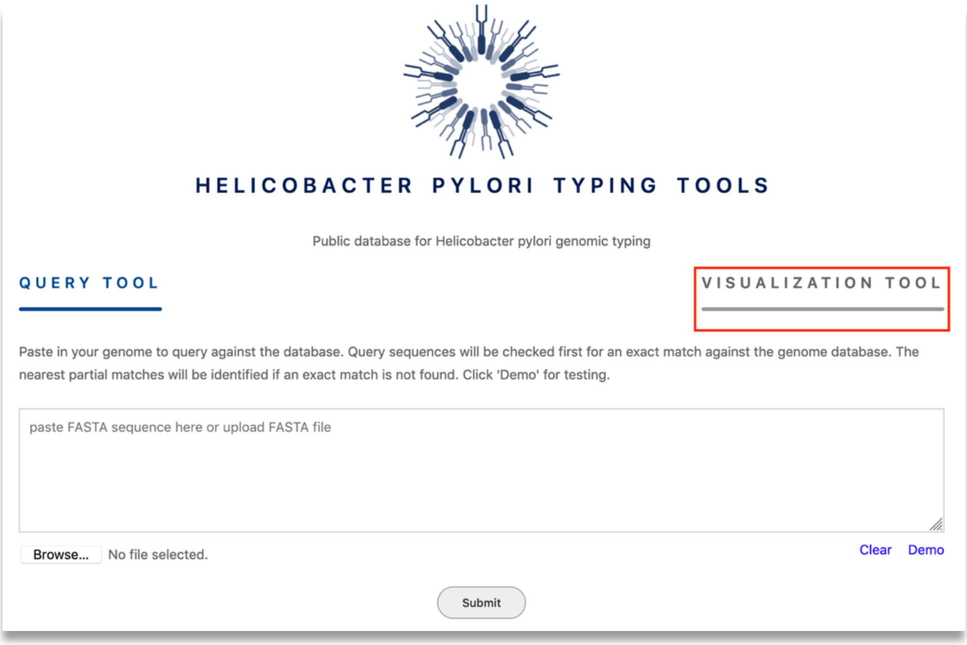

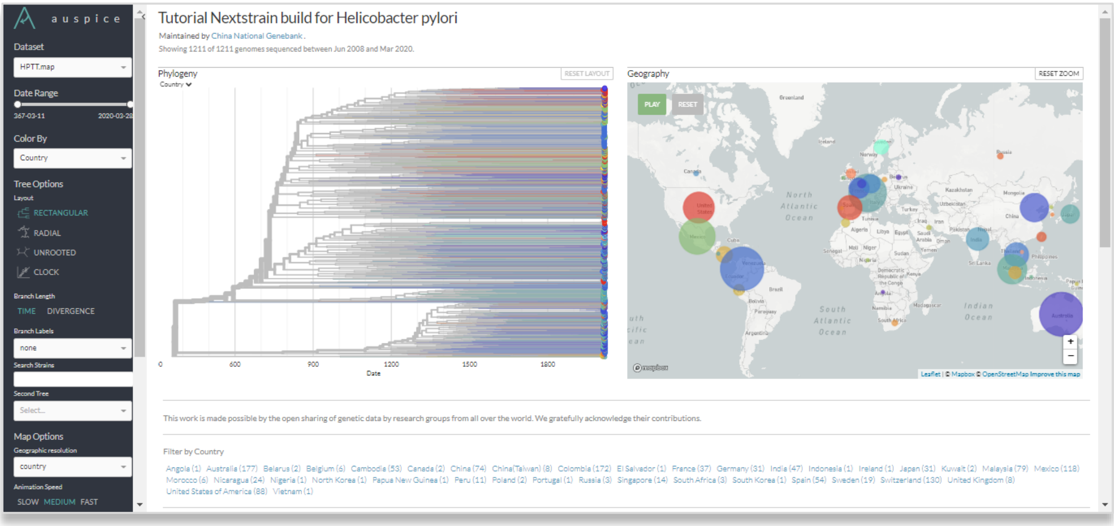

#### - The linking functions

When the genome matched the data in our database, it is easily to see the geographic location by clicking ID.

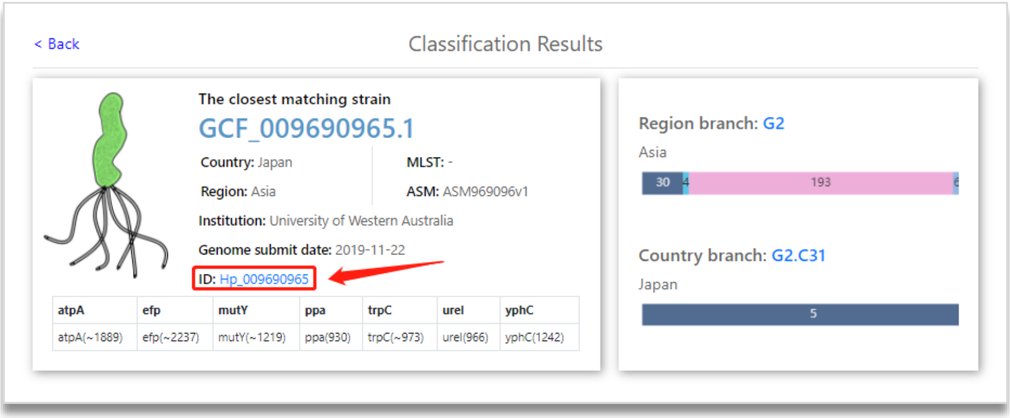

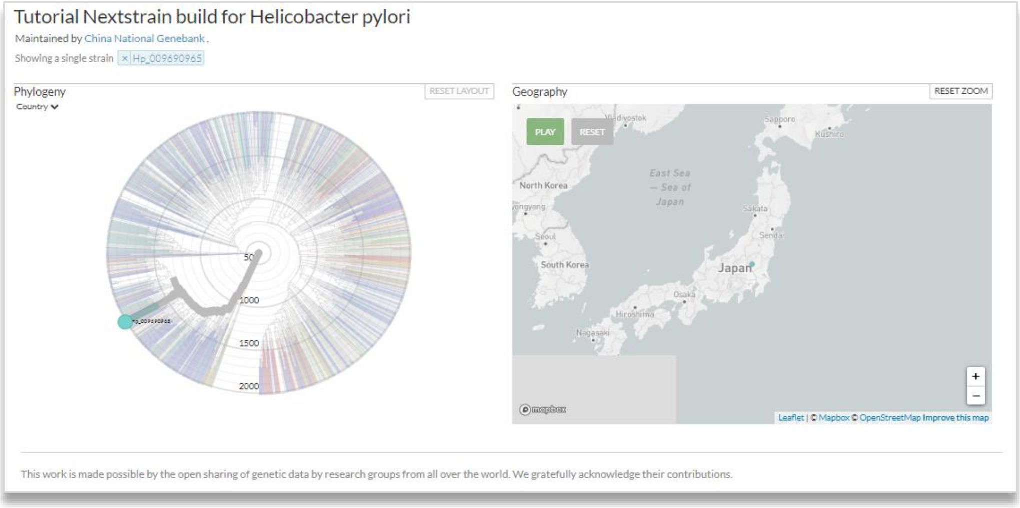

When the genome matched the data in our database, it is easily to see the phylogenetic relationship and geographic location by clicking the group ID (the letters in blue).

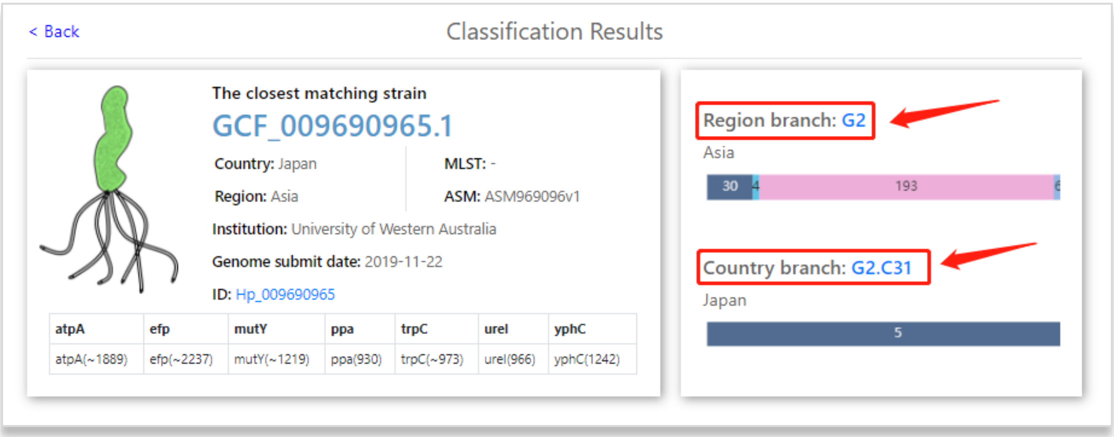

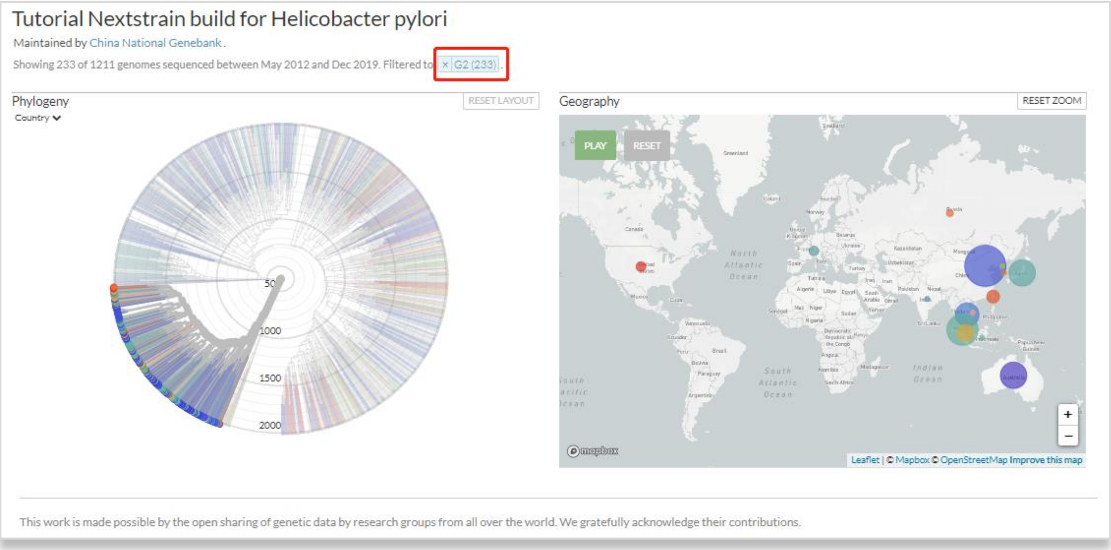
Tutorial Nextstrain build for Helicobacter pylori
Maintained by China National Genebank.
Showing 233 of 1211 genomes sequenced between May 2012 and Dec 2019. Filtered to × G2 (233).
Phylogeny
Country
RESET LAYOUT
500
1000
1500
2000
Geography
RESET ZOOM
PLAY
RESET
Leaflet | © Mapbox © OpenStreetMap Improve this map
This work is made possible by the open sharing of genetic data by research groups from all over the world. We gratefully acknowledge their contributions.

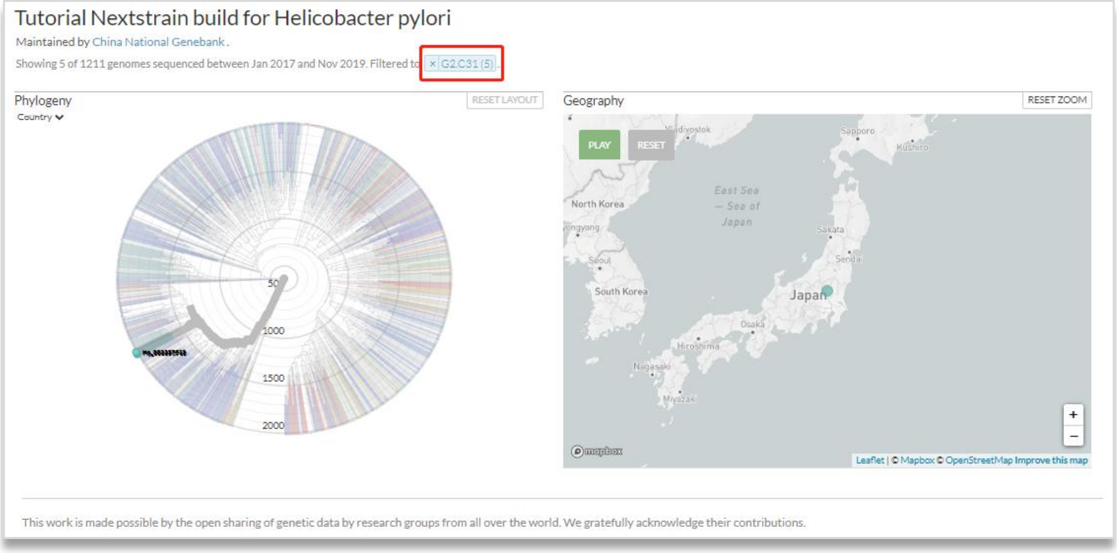
Tutorial Nextstrain build for Helicobacter pylori
Maintained by China National Genebank.
Showing 5 of 1211 genomes sequenced between Jan 2017 and Nov 2019. Filtered to × G2.C31 (5).
Phylogeny
Country
RESET LAYOUT
500
1000
1500
2000
Geography
RESET ZOOM
PLAY
RESET
North Korea
South Korea
Japan
East Sea — Sea of Japan
Seoul
Osaka
Hiroshima
Nagasaki
Miyazaki
Sapporo
Kushiro
Sakata
Sendai
Leaflet | © Mapbox © OpenStreetMap Improve this map
This work is made possible by the open sharing of genetic data by research groups from all over the world. We gratefully acknowledge their contributions.
